# Intrinsic DNA topology as a prioritization metric in genomic fine-mapping studies

**DOI:** 10.1101/837245

**Authors:** Hannah C. Ainsworth, Timothy D. Howard, Carl D. Langefeld

**Affiliations:** Department of Biostatistics and Data Science, Wake Forest School of Medicine, Winston-Salem, NC, 27157, USA; Department of Biochemistry, Wake Forest School of Medicine, Winston-Salem, NC, 27157, USA

## Abstract

In genomic fine-mapping studies, some approaches leverage annotation data to prioritize likely functional polymorphisms. However, existing annotation sources often present challenges as many: lack data for novel variants, offer no context for noncoding regions, and/or are confounded with linkage disequilibrium. We propose a novel annotation source – sequence-dependent DNA topology – as a prioritization metric for fine-mapping. DNA topology and function are well-intertwined, and as an intrinsic DNA property, it is readily applicable to any genomic region. Here, we constructed and applied, Minor Groove Width (MGW), as a prioritization metric. Using an established MGW-prediction method, we generated an MGW census for 199,038,197 SNPs across the human genome. Summarizing a SNP’s change in MGW (ΔMGW) as a Euclidean distance, ΔMGW exhibited a strongly right-skewed distribution, highlighting the infrequency of SNPs that generate dissimilar shape profiles. We hypothesized that phenotypically-associated SNPs can be prioritized by ΔMGW. We applied Bayesian and frequentist MGW-prioritization approaches to three non-coding regions associated with System Lupus Erythematosus in multiple ancestries. In two regions, including ΔMGW resolved the association to a single, trans-ancestral, SNP, corroborated by external functional data. Together, this study presents the first usage of sequence-dependent DNA topology as a prioritization metric in genomic association studies.

**Graphical Abstract:** We hypothesize that SNPs imposing dissimilar minor groove width profiles (ΔMGW) are more likely to alter function. ΔMGW was interrogated genome-wide and then used as a weighting metric for fine-mapping associations.

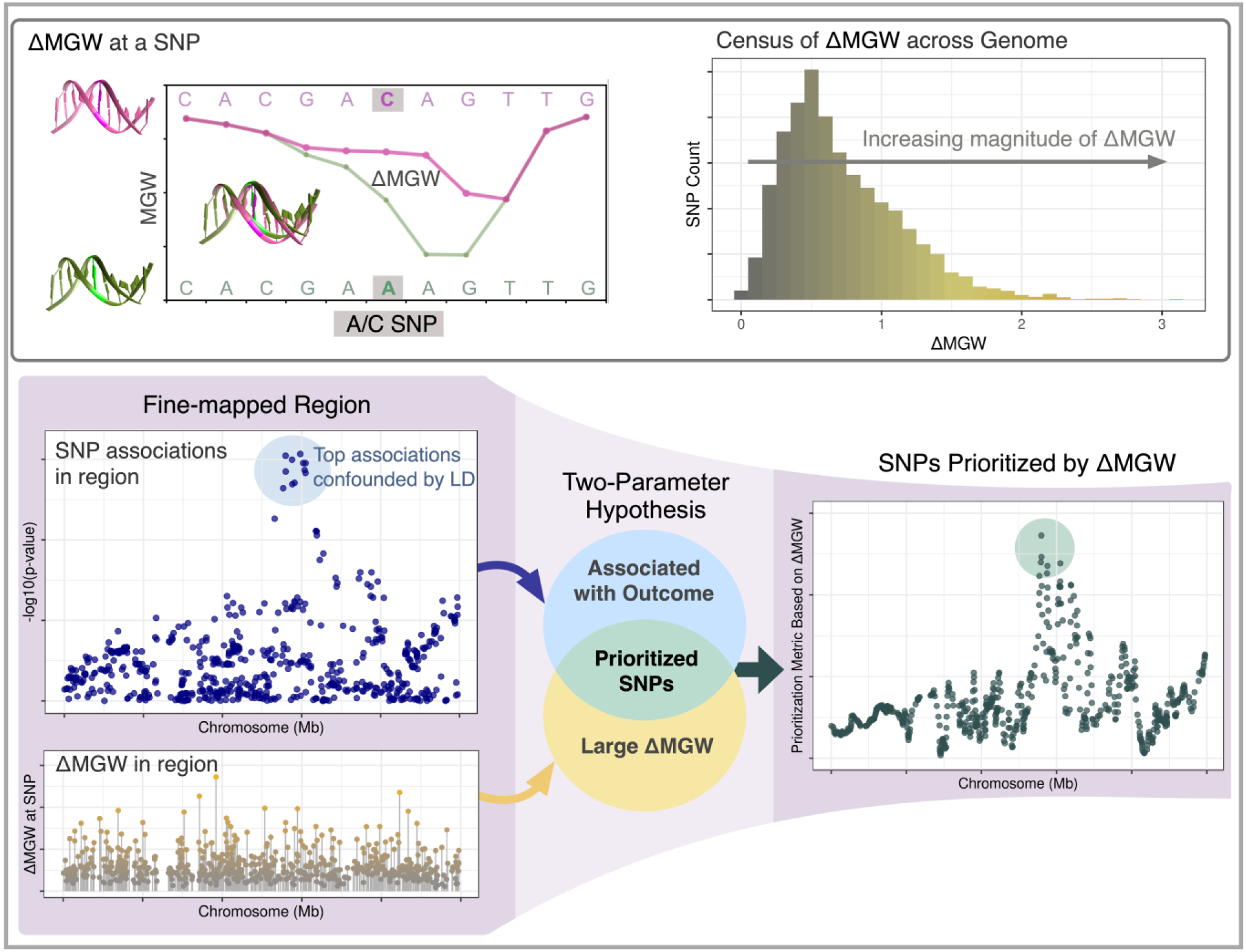

## Introduction

Genetic association studies have successfully identified thousands of loci associated with a broad range of phenotypes.(1) However, despite the abundance of these genomic associations, analytic challenges have largely hindered identification of the specific genomic drivers of disease.(2–4) First, linkage disequilibrium (LD) constitutes a major analytic challenge, as highly correlated variants exhibit comparable evidence of association, making it difficult to statistically isolate causal polymorphisms. Second, many associated single nucleotide polymorphisms (SNPs) reside in non-coding regions, occluding functional relevance without additional context and information. Even with increased sample sizes and variant coverage, these challenges remain.(2–5) In-depth functional analyses are not practical for a large number of variants, and thus, there remains the need to effectively prioritize the most likely causal variants for follow-up studies and approaches (e.g. CRISPR).

To prioritize potential causal variants, association results can be weighted by external functional information (e.g. histone modifications, eQTL status, transcription factor binding sites).(5–8) This approach has been successful in reducing and refining associated variants, and there are a growing number of tools and methods that integrate external data with genomic association studies.(6, 9–13) Nevertheless, such methods are not without limitations. Importantly, the choice of annotation and database bias are strong factors for consideration as missing or incomplete functional data could result in down-weighting potentially causal polymorphisms. These challenges particularly arise for regions with no (presently) known functional implications. Additionally, many annotation resources are based on European data; and thus may offer limited information for genetic studies in non-European individuals (e.g. novel regions).(14, 15) Such limitations can reduce the rate of progress in understanding the functional impact of ancestry-specific associations and perpetuate health disparities.(16, 17) To alleviate some of these biases imposed by external datasets, we propose a prioritization approach that leverages information intrinsic to the DNA itself, sequence-dependent DNA topology.

From chromatin conformation to selective protein binding,(18–26) DNA is a highly dynamic macromolecule with structure inherently linked to function. Sequence-dependent DNA topology (or shape) refers to the geometric parameters (measured in Angstroms or degrees) between successive nucleotides in a DNA sequence.(24, 27–29) The sequence dependency of these spatial measures (**Figure 1**) has been well-studied and in recent years, increasingly connected to various functional implications, including protein binding, DNA stability, and methylation.(18, 20, 21, 23, 30–38) High-throughput DNA shape prediction methods now enable exploration of DNA topology on a genome-wide scale, and thus, provide new opportunities in association studies.(24, 39)

**Figure 1:**
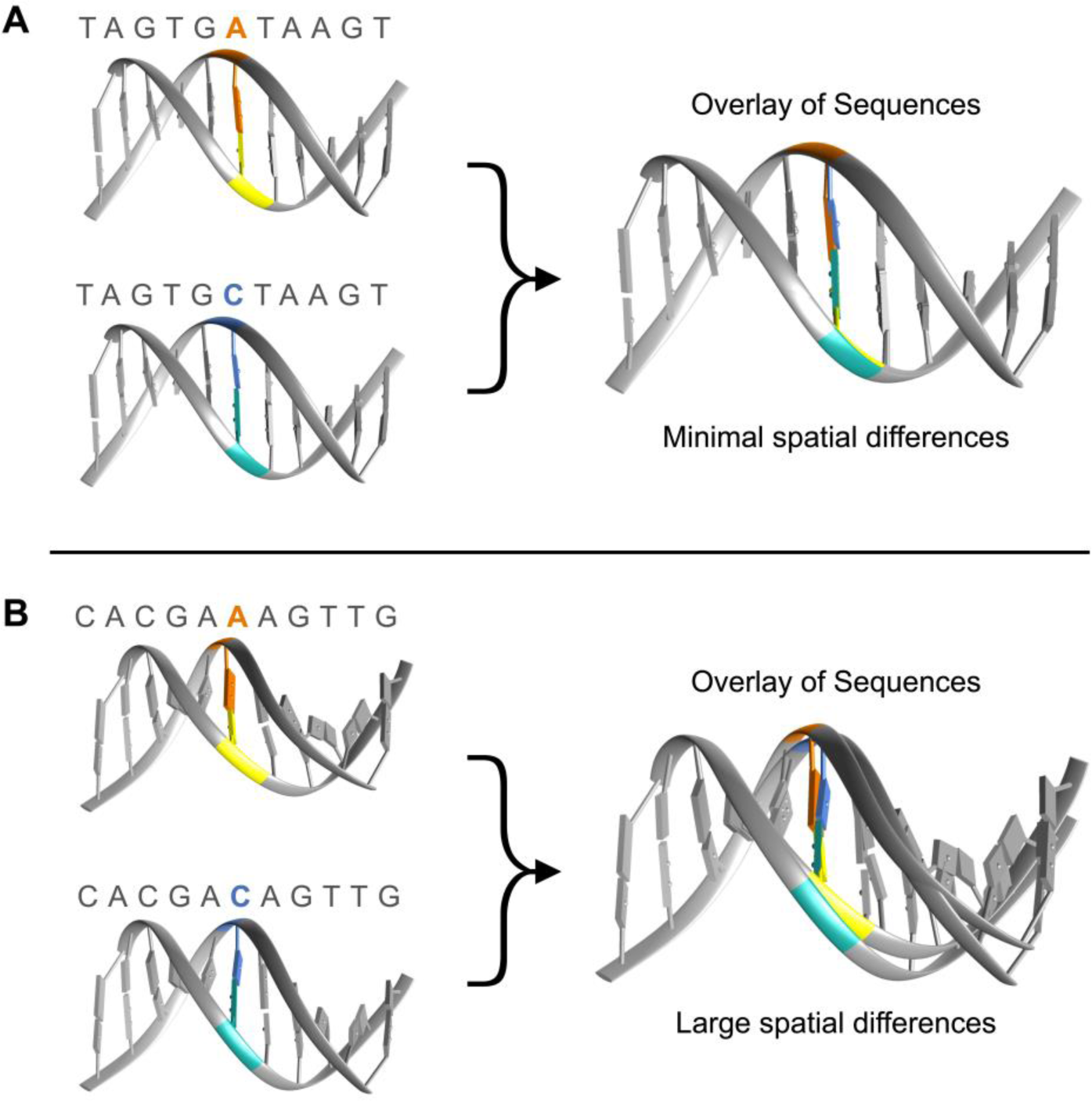
Single nucleotide substitutions sequence can impose large or small changes on local DNA shape, dependent on the flanking sequence. (A) A single A/C substitution within a sequence generates minimal spatial differences. (B) A single A/C substitution within a sequence imposes large spatial differences

This study presents using sequence-dependent DNA topology as a prioritization metric in genomic association studies. Here, we focused on minor groove width (MGW), which measures the distance (Angstroms, Å) between the sugar phosphate backbone of the forward and reverse strands. For each SNP, we analyzed its change in minor groove width (ΔMGW) to evaluate whether the SNP’s alleles created similar or divergent MGW profiles. MGW has been implicated in numerous protein binding studies and used in transcription factor binding prediction algorithms.(18, 20, 24, 32, 34, 36, 37, 40, 41) Recently it was studied in the context of purifying selection, where “shape disrupting variants” (examples shown in **Figures 2** and **3**) tend to be less common in functional regions (shape-preserving polymorphisms being more frequent).(42) Thus, we proposed that if a phenotypically-associated SNP also yields a large ΔMGW, it is more likely to be causal as a function of divergent shape profiles.

**Figure 2:**
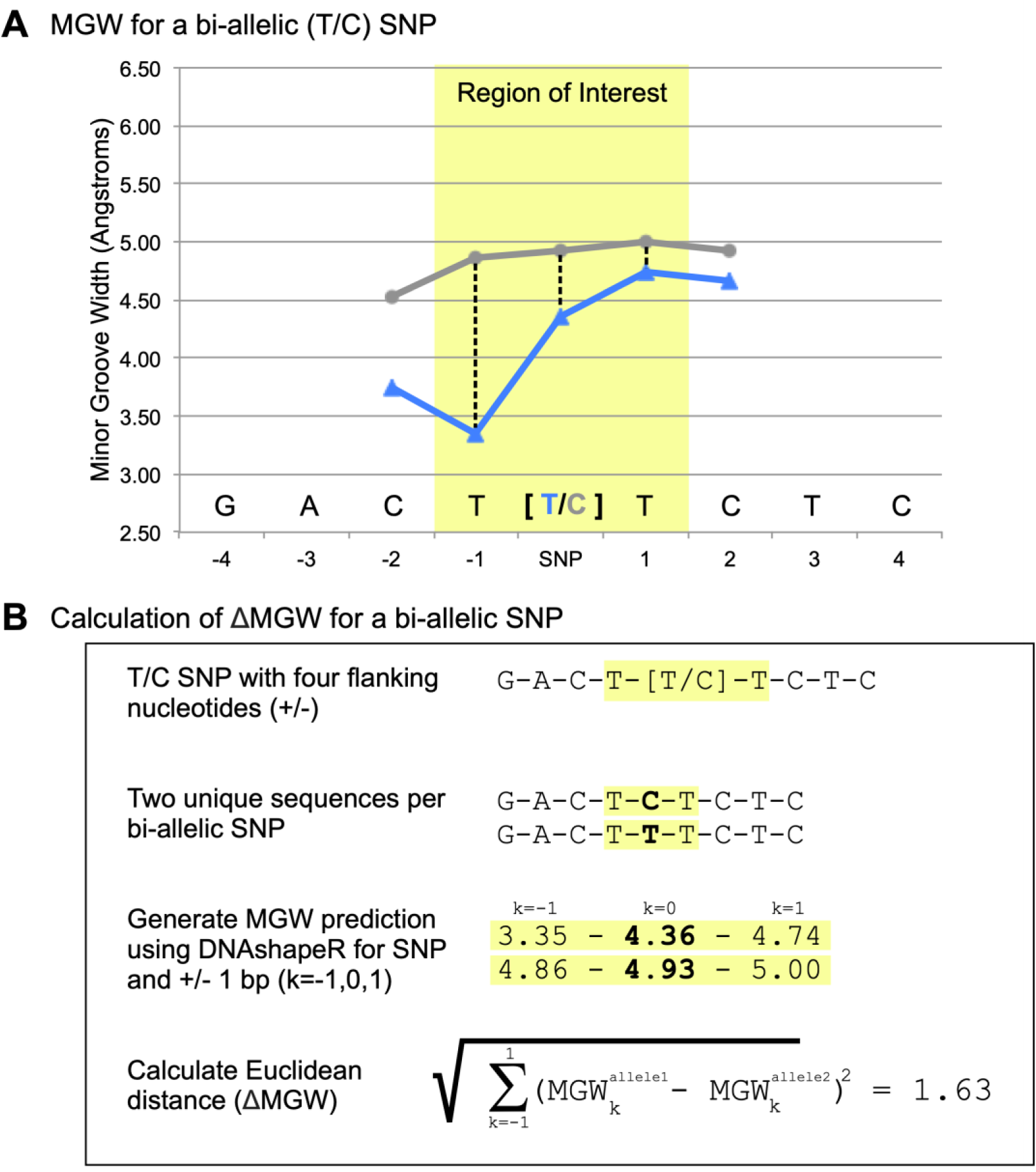
Generation of ΔMGW for a SNP. (A) Minor groove width measures are plotted for the two sequences generated by a specific bi-allelic T/C SNP. For a given SNP, the flanking sequence (+/− 4 bp) was used as input for DNAshapeR (via Bioconductor) which calculates MGW along a rolling sequence window. For a 9-mer sequence, the MGW can be consistently provided at the SNP’s position +/− one nucleotide which is highlighted in yellow and labeled as the ‘region of interest’. Expanding this region to additional nucleotides would require a longer input sequence and increases chance of additional variants being within the input (and introducing additional variability). Although the two sequences for a SNP only differ at one nucleotide (at the SNP position), the impact on MGW carries through adjacent bases. Thus, ΔMGW was calculated to capture the change in MGW for a SNP by incorporating information at the SNP’s position and +/− 1 base pair (dashed lines). (B) Workflow for calculating the ΔMGW for a bi-allelic SNP. This method captures the change in MGW at the SNP position and +/− 1 base pair. This Euclidean distance captures ΔMGW as a measure of magnitude (in Angstroms).

**Figure 3:**
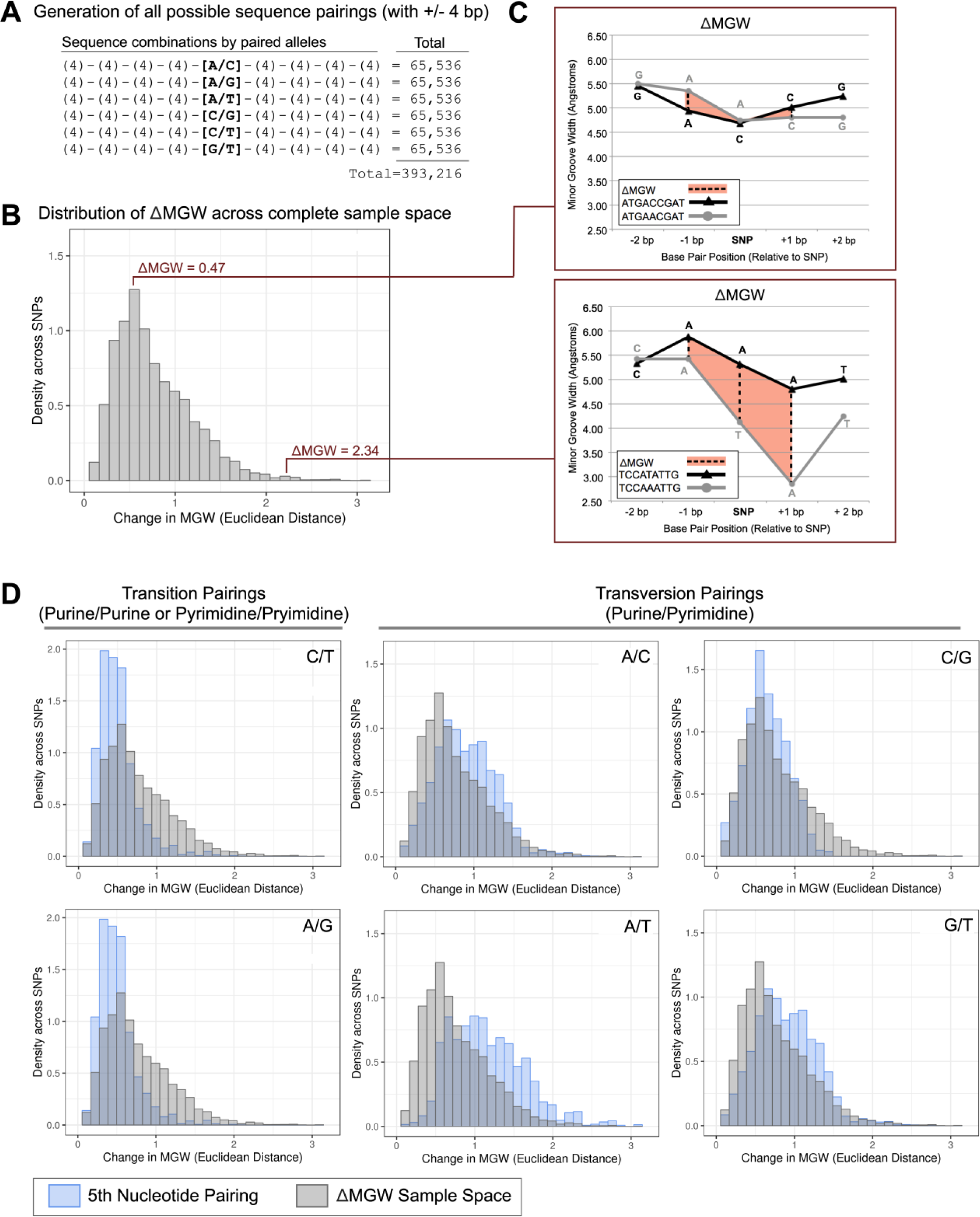
Summarization of ΔMGW across the complete sample space. (A) ΔMGW sample space was constructed on six allele pairings (A/C, A/G, A/T, C/G, C/T, G/T) with all possible combinations for flanking +/− 4 bp. This yielded 393,216 paired sequences that were evaluated for ΔMGW. (B) The distribution of ΔMGW for the 393,216 paired sequences, these summary statistics are listed in Table 1. (C) Two randomly selected paired sequences from the average and right tail of the ΔMGW distribution are shown. Sequences are plotted with their respective MGW values (Angstroms). ΔMGW is calculated as a Euclidean distance, which captures the change in MGW (dashed lines) at the SNP position and +/− 1bp (highlighted in orange). ATGA[C/A]CGAT exhibits a small ΔMGW, at 0.47 Å while TCCA[T/A]ATTG yields a large change in MGW (2.34 Å) which we would hypothesize to have greater potential for functional consequence if also associated with disease status. (D) The ΔMGW distribution for all paired sequences (gray) is shown superimposed on the ΔMGW distributions by 5th nucleotide alleles (blue). Transition pairings (C/T, A/G) have a more strongly skewed distribution with a smaller average ΔMGW compared to transversion pairings (A/C, A/T, C/G, G/T), (Table 1). Pairings that represent complimentary sequences (C/T – A/G and A/C – T/G) exhibit the same distributions of ΔMGW, as expected.

We specifically hypothesized that highly correlated SNPs in a phenotype-associated region can be functionally prioritized using each SNP’s magnitude of ΔMGW. We evaluated this hypothesis in three stages. First, using an established MGW-prediction algorithm(39), we generated the complete sample space for ΔMGW for all possible input sequences. Second, we evaluated the observed frequency of ΔMGW across the human genome using bi-allelic SNPs in the dbSNP SNP150 dataset. Third, we tested this approach by prioritizing SNPs in three genomic regions previously associated with systemic lupus erythematosus (SLE)(43) leveraging both frequentist and Bayesian association methods.

## Methods and Materials

### Calculation of ΔMGW for a bi-allelic SNP

The predicted MGW for a given sequence was obtained using the DNAshapeR package (https://bioconductor.org/packages/release/bioc/html/DNAshapeR.html), available through Bioconductor.(39) DNAshapeR calculates DNA features using Monte Carlo simulations for nucleotide structure based on DNA sequence fragments. DNA feature predictions are based on a rolling window of five nucleotides for a given n-length sequence. For this study, to capture the MGW at a SNP, we used the four flanking (up and downstream) nucleotides (9-mer sequence) as input. Each bi-allelic SNP produces two unique 9-mer sequences (one sequence for each allele) and thus, both of a SNP’s sequences were submitted to DNAshapeR to obtain the corresponding feature vectors for MGW. The MGW was retained for the nucleotide at the SNP’s position as well as +/− 1 nucleotides. Capturing MGW for additional bases would require longer input sequences, which could introduce additional variability (e.g. SNPs within the flanking sequence). The ΔMGW was calculated as a Euclidean distance for the SNP and +/− 1 base (**Figure 2**).

### Generation of ΔMGW sample space

To calculate the entire sample space for ΔMGW, we generated a dataset of all possible input sequences. Since our goal was to evaluate the ΔMGW at a SNP with +/− 4 base pairs, input sequences required nine nucleotides. Thus, all possible combinations of Adenine, Cytosine, Guanine, and Thymine, generated 262,144 9-mer sequences. From this dataset, all possible bi-allelic pairings (A/C, A/G, A/T, C/G, C/T, G/T) were created on the 5^th^ nucleotide of each sequence (“SNP position”) while holding the flanking nucleotides constant, generating 393,216 9-mer pairings. These 9-mer pairings represent every possible sequence combination that could be observed for a bi-allelic SNP (**Figure 3**). These paired sequences were evaluated for ΔMGW using the previously described method.

### Visualization of DNA sequences

DNA shape measures, provided by DNAshapeR, were submitted as a parameter file to the 3D-Dart webportal (http://milou.science.uu.nl/services/3DDART/) for a ‘BDNA nucleic acid’.(44) Resulting pdb files from 3D-Dart were then visualized using Chimera (https://www.cgl.ucsf.edu/chimera/).(45)

### Curating dbSNPs150 database

The NCBI hg19 dbSNPs150 data file (snp150.txt.gz) was downloaded via UCSC GoldenPath (hgdownload.cse.ucsc.edu) on July 6, 2018.(46) Insertion-deletions, tri-allelic, quad-allelic, and multiple nucleotide polymorphisms were excluded. Retained bi-allelic SNPs were limited to those located on chromosomes 1-22 and X. Any SNPs that were labeled with “Unusual Conditions” as defined by UCSC were excluded, as these indicate possible discrepancies among alleles and/or potential mapping issues (e.g. SNP flanking sequence aligns to more than one location in the reference assembly).(46, 47) The pruned bi-allelic dataset contained 199,038,272 SNPs.

For dbSNP 150 data, each SNP’s flanking sequence of four nucleotides was retrieved from the Human Reference Genome (downloaded October 2017)(48) using SAMTOOLS. For each SNP, the dbSNP “Strand” variable was used to inform if the alleles reported by dbSNP aligned with the reference genome. All SNPs were successfully queried against the reference genome. There were 75 SNPs that contained at least one flanking base encoded as “N” (any base) and were excluded from summarizations, leaving a final dataset of 199,038,197 SNPs. The ΔMGW for these sequences were obtained as described above.

### SLE Immunochip Data for fine-mapping analyses

Genomic data for fine-mapping analyses came from the published trans-ancestral SLE Immunochip study; genotype calling and genomic quality control methods were previously described.(43) This data includes three ancestries, European Ancestry (EA), African Ancestry (AA), and Hispanic Ancestry (HA), with large case-control counts: EA (6,748; 11,516), AA (2,970; 2,452), and HA (1,872; 2,016).

Genomic regions were named for the genes in physical proximity to the region of association. Non-HLA genomic regions were selected for fine-mapping if the region contained SNPs reaching genome-wide significance (p< 5×10^−8^) in at least two ancestry-specific analyses.(43) We also limited our analyses to regions where the top associations mapped to non-coding regions (e.g. introns, intergeneic), where we hypothesize DNA topology might provide novel insight to the fine-mapping analyses. Genomic regions containing *FAM167A-BLK* (8p23), *STAT4* (2q32), and *TNIP1* (5q33) met these criteria. Quality controlled genomic data for these regions were extracted using a 250 kb window around the previously reported top association from the Immunochip analysis.(43)

SNPs from the selected genomic regions were queried against the human reference genome to retrieve the four flanking bases. Each SNP’s strand information (based on Illumina Infinium Immunochip documentation) was utilized to ensure that the corresponding alleles appropriately aligned with the reference genome.

### Statistical Analyses

#### Single-SNP associations

Single-SNP associations were previously reported and described in the transancestral SLE Immunochip study.(43)

#### SKAT analyses

The previous single-SNP logistic regression analyses (43) did not incorporate SNP-specific weights/information. Thus, SNPs in high LD yielded comparable association values. The Sequence Kernel Association Test (SKAT) is a regression approach that was designed to handle covariates and SNP-specific weights through a weighted linear kernel.(49) It was shown that well-selected SNP weights can yield better statistical power (e.g. increasing weight of functional variants).(49) SKAT was originally developed to leverage minor allele frequency (MAF), as the weighting scheme in rare variant studies; however, the SKAT framework is a general method that can accommodate any user-specified SNP weights.(49) Here, we used ΔMGW as the weighting scheme. A variation of SKAT is the Optimal unified test which combines both SKAT and the burden test (SKAT-O).(12) The SKAT-O test statistic is a weighted average of the SKAT and burden test statistics and can be beneficial when applying to genomic regions where one test may be better powered than another.(50) Primary advantages of burden tests occur when a large number of variants are causal and for smaller sample sizes (SKAT loses power in small sample sizes, <2000 cases and controls). Generally, burden tests do not perform as well as SKAT when a large proportion of the variants are non-causal.(12, 49, 50) In this study, our datasets are large (AA: 5,422; EA: 18,264; HA: 2,016), and we expect many of the highly associated SNPs in LD to be non-causal; thus, in this scenario we selected SKAT to be more appropriate, which is consistent with published power calculations and simulations.(12, 49, 50) SKAT was applied to genomic regions through its implementation in the R package, SKAT (https://CRAN.R-project.org/package=SKAT). For each genomic region, the model parameters and residuals were calculated for SKAT using SKAT_Null_Model() for a dichotomous outcome (case/controls status) and previously described (43) population-specific factors (to account for admixture). Since all datasets (AA, EA, and HA) had a sample size greater than 2,000 cases and controls, no small-sample adjustment was applied. Within each genomic region, adjacent 5-SNP windows were generated, offset by 1 SNP. Each window was evaluated using the SKATbinary() with method=SKAT and a linear-weighted kernel with SNPs weighted by their ΔMGW. To evaluate consistency of the results (e.g. for SNPs outside of the main peak of association), genomic regions were also evaluated using equal-weighting for all SNPs. Given the small window size (n=5 SNPs), we expect a large proportion of each window to contain non-causal SNPs, further supporting our selection of SKAT. For comparison, we also applied SKAT-O but noted minimal differences on the final outcome. To localize the top association signals to each SNP, SNP-window p-values were treated as a SNP prioritization metric by generating the geometric mean of −log_10_(p-values) across windows containing each SNP. That is, the prioritization metric was calculated using the p-value for each SKAT analysis window (p_i_) that contained the *k^th^* SNP (*n* analysis windows). With the exception of the first and last five SNPs in a region, each SNP_k_ was included in five analysis windows (n=5). Thus, for each SNP *k*, we calculated its prioritization metric as:

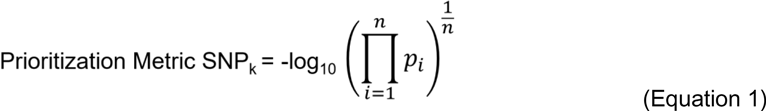

#### Bayesian Approach: Credible SNP Sets

Frequentist approaches, such as those implemented SKAT or single-SNP logistic regression analyses are widely utilized; however, their resulting p-values are not without limitations.(51) For one, p-values do not capture the confidence of a particular association. Furthermore, they’re more dependent on factors such as the power of the statistical test (influenced by sample size and other variables). Bayesian methods offer an alternative approach; here, Bayes factors are used, capturing the ratio of probabilities between the null and alternative hypotheses.

As a comparison to the frequentist approaches, we used SNPTEST to generate the Bayes factors (BF), using the score test and additive genotype modeling.(52) Posterior probabilities for a given SNP *k*, were then calculated using method published by the Welcome Trust Case Control Consortium.(53) For SNPs 1-j in the region, the posterior probability for each SNP *k*, was calculated by:

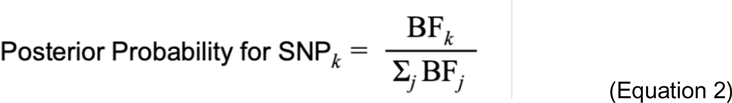

Using these posterior probabilities, the 95% credible set was determined for each region. This test assumes only one causal SNP in the region and places equal *a priori* probabilities that the causal SNP is any one of the analyzed SNPs.(53) In this study, we applied this method to previously defined regions (43) where we hypothesized the association signal is driven by one SNP.

Like the single-SNP logistic regression analyses, this Bayesian analysis is not weighted by functional data. Thus, for a ΔMGW-weighted analysis, a derived credible set was generated from posterior probabilities that accounted for each SNP’s ΔMGW through *ad hoc* weighting, where the posterior probability for a given SNP *k*, was calculated by weighting the Bayes factor by ΔMGW_k_ divided by the weighted average of Bayes factors for SNPs 1-j in the region. Here, the derived posterior probability for each SNP *k*, is:

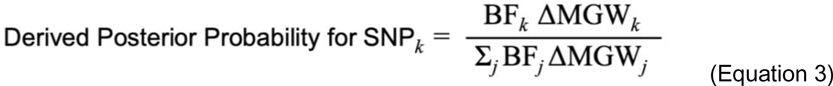

Using these values, the derived 95% credible SNP sets were generated and compared with the unweighted 95% credible SNP sets. This methodology enabled weighting by a continuous variable versus existing methods designed for dichotomous (presence/absence of functional annotation) SNP weights.(54)

### Functional Annotation

To evaluate the functional plausibility for an identified variant, several publically available resources were referenced. For variant associations with gene expression (eQTL status), the Genotype-Tissue Expression (GTEx) dataset, version 7 (hg19) was queried at gtexportal.org.(55) GTEx is a comprehensive eQTL resource, providing eQTL information across 48 tissues. SNPs were also queried using the SCREEN (Search Candidate cis-Regulatory Elements by Encode, http://screen.encodeproject.org).(56, 57) Built using Encode data, SCREEN (hg19) evaluates if a given genomic coordinate resides in a Candidate cis-Regulatory Element (ccRE). ccREs are designated based on evidence from DNase hypersensitivity sites, H3K4me3 and H3K27ac histone activity, and CTCF-binding data. SCREEN contains 1.31 million ccREs, correlating to 20.8% of the mappable human genome (http://screen.encodeproject.org). Genomic variants were also evaluated for evidence of long-range DNA interaction via Hi-C data (hg19) available through the Yue Lab 3D Genome Browser (http://promoter.bx.psu.edu/hi-c/).(58) Similar to the ccRE search, SNPs were queried to see if they resided in a genome region that exhibited long-range chromatin interactions. The Yue Lab’s Capture Hi-C data offers information across 19 cell line options. We evaluated immune-related cell types: naïve B-Cells, CD4_Total (CD4 activated and Naïve), CD8 naïve, monocytes, and neutrophils.

## Results

### For ΔMGW, SNPs in the human genome exhibit a stronger right skewed distribution in comparison to the complete sample space

In the complete sample space of ΔMGW, ΔMGW values ranged from 0.00 to 3.16 Å, with a mean of 0.77 Å and a standard deviation of 0.42. **(Table 1)** The overall data exhibited a right-skewed distribution (**Figure 3**) with few sequences inducing large changes in MGW. Unsurprisingly, given the sequence-dependency of this topological measure, parsing the data by the paired alleles (fifth nucleotide, see Methods and Materials), revealed allele-specific patterns of ΔMGW **(Table 1)**. Transition pairings (A/G and C/T) yielded the smallest changes in ΔMGW, while transversion pairings (Purine/Pyrimidine) produced the largest changes in ΔMGW. Subsets that represent complimentary allele pairs (i.e. A/G & T/C; A/C & T/G) yielded the same ΔMGW values. (**Table 1)** Of all allele-pairings, A/T alleles presented the largest ΔMGW with a mean of 1.16 Å (SD, 0.47) **(Figure 3)**.

**Table 1.** Summary statistics for the complete ΔMGW (Å) sample space.

| 5<sup>th</sup> Nucleotide pairing<sup>a</sup> | N | Min. | Max. | Range | Median | Mean | Standard Deviation |
| --- | --- | --- | --- | --- | --- | --- | --- |
| A/C | 65,536 | 0.03 | 2.74 | 2.71 | 0.86 | 0.90 | 0.39 |
| A/G | 65,536 | 0.05 | 2.07 | 2.02 | 0.46 | 0.50 | 0.25 |
| A/T | 65,536 | 0.07 | 3.16 | 3.09 | 1.11 | 1.16 | 0.48 |
| C/G | 65,536 | 0.00 | 1.44 | 1.44 | 0.62 | 0.64 | 0.27 |
| C/T | 65,536 | 0.05 | 2.07 | 2.02 | 0.46 | 0.50 | 0.25 |
| G/T | 65,536 | 0.03 | 2.74 | 2.71 | 0.86 | 0.90 | 0.39 |
| All Possible | 393,216 | 0.00 | 3.16 | 3.16 | 0.67 | 0.77 | 0.42 |

<sup>a</sup>Pairings generated by 5<sup>th</sup> nucleotide in 9-mer sequence, all other nucleotides held constant.

We compared the ΔMGW sample space statistics to the observed frequencies of ΔMGW across the human genome using dbSNP data. The hg19 download of NCBI dbSNP150 contained 234,104,110 entries. After pruning to high quality (see Methods and Materials), bi-allelic SNPs, 199,038,197 polymorphisms remained. For these SNPs, there was an average ΔMGW of 0.68 Å with a standard deviation of 0.43. In comparison to the ΔMGW sample space, SNPs across the genome exhibited a stronger, right-skewed distribution of ΔMGW. (**Figure 3**). Transition SNPs are more likely to occur (59, 60), and this is consistent with our SNP150 summarizations, where transition SNPs comprised 66.43% of the dataset (**Table S1**). Our ΔMGW sample space summarization showed that transition allele pairings had the smallest change in ΔMGW **(Table 1);** thus, the decreased average in ΔMGW dbSNP data is expected and illustrates the high prevalence shape-preserving SNPs in the genome. To evaluate patterns in ΔMGW by SNP function (i.e. missense, intron, coding-synonymous), SNPs with a single NCBI-designation (see Methods and Materials) were subset and summarized (**Table 2, Figure 4**). Notably, some SNP categories are limited to specific sequence combinations(61) (i.e. stop-loss, **Table S2**), which were reflected in the SNP-function-specific patterns of ΔMGW. (**Figure 4**) Coding-synonymous SNPs exhibited the smallest overall change in ΔMGW (mean=0.48 Å). Unknown and intron SNPs, which are not constrained to specific sequences (by definition), comprised the two largest categories (n_unknown_=99,004,130; n_intron_=84,909,115) and yielded high averages for ΔMGW: 0.69 Å and 0.56 Å, respectively.

**Figure 4:**
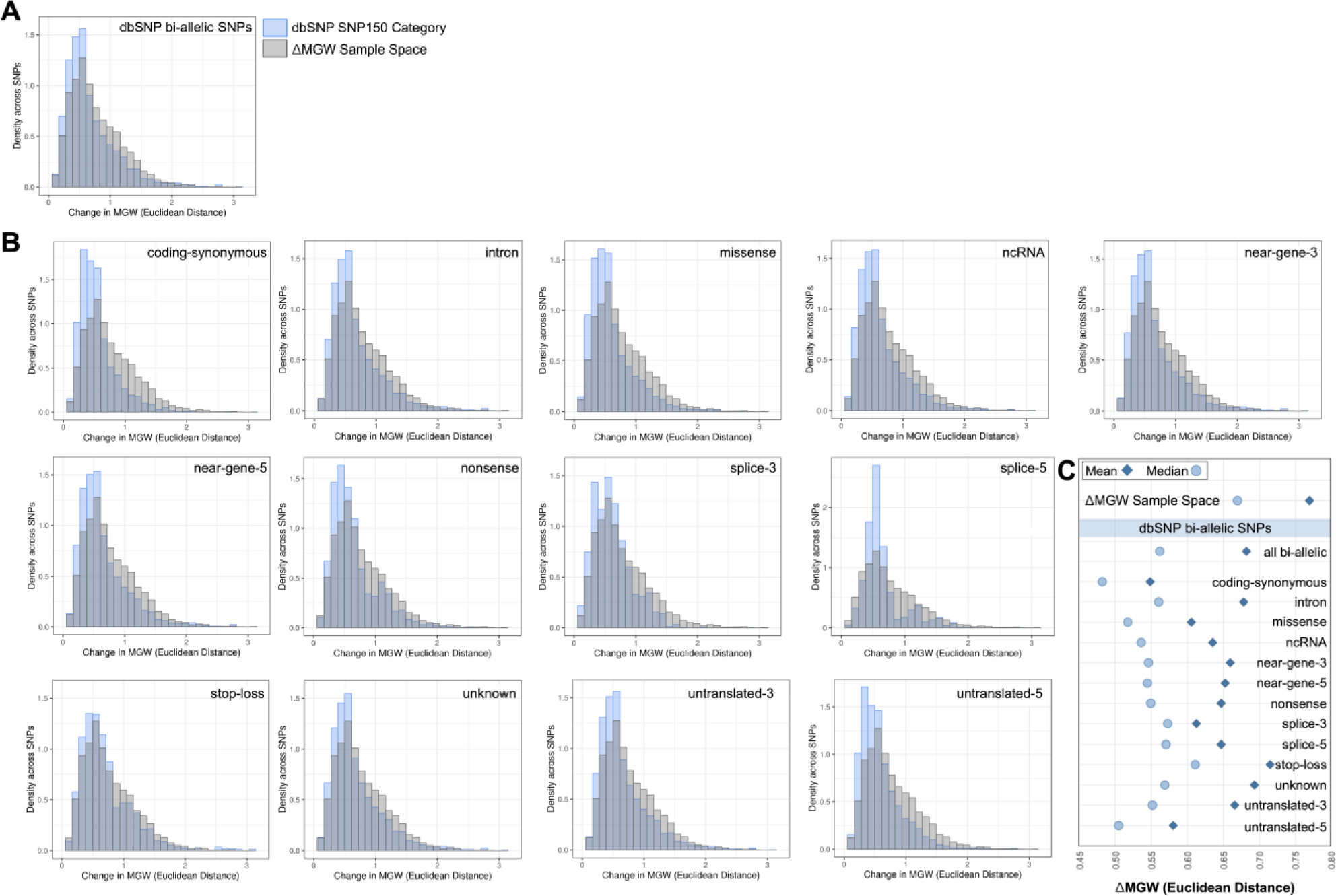
Summarization of ΔMGW across the human genome using bi-allelic SNPs from dbSNP SNP150. (A) Comparison of ΔMGW sample space (Figure 3) and the observed ΔMGW from SNPs across the genome (via dbSNP). Distribution of ΔMGW is shown in blue for observed bi-allelic SNPs from the SNP150 dataset (n=199,038,197 SNPs). The ΔMGW sample space distribution (Figure 3) is plotted in gray (n=393,216 paired sequences). The observed ΔMGW across genomic SNPs showed a stronger right skewed distribution than what would be expected from a random sampling of the entire sample space of all-possible sequences. Only small numbers of SNPs elicit large magnitudes of ΔMGW. (B) ΔMGW distributions are similarly shown for SNP subsets, by NCBI function (exclusive NCBI function label for each SNP, see Methods and Materials). Again, each distribution is superimposed with the distribution from the ΔMGW sample space (shown in gray). Subsetting by NCBI function yields similar patterns observed in part A, with observed genomic SNPs showing smaller averages in ΔMGW. Some NCBI SNP-functions have specific sequence requirements (Supplemental Table 1) and these are reflected in the resulting ΔMGW distributions which are also sequence-dependent (e.g. splice-6, nonsense). (C) The mean and median ΔMGW for each SNP category. All dbSNP SNP categories have significantly lower mean and median compared to the ΔMGW sample space (Tables 1-2). Coding-synonymous SNPs have the smallest magnitudes of ΔMGW, compared to all other categories.

**Table 2.** Summary Statistics for ΔMGW (Å) across bi-allelic SNPs in dbSNP SNP150 dataset.

| SNP Category |  | N | Min. | Max. | Range | Median | Mean | Standard Deviation |
| --- | --- | --- | --- | --- | --- | --- | --- | --- |
| dbSNP SNP150 (bi-allelic) |  | 199,038,197 | 0.00 | 3.16 | 3.16 | 0.56 | 0.68 | 0.43 |
| Single-NCBI Function Subsets | coding-synonymous | 1,178,980 | 0.00 | 2.58 | 2.58 | 0.48 | 0.55 | 0.30 |
|  | intron | 84,909,115 | 0.00 | 3.16 | 3.16 | 0.56 | 0.68 | 0.42 |
|  | missense | 2,345,831 | 0.00 | 3.16 | 3.16 | 0.52 | 0.61 | 0.36 |
|  | ncRNA | 499,593 | 0.00 | 3.16 | 3.16 | 0.54 | 0.63 | 0.38 |
|  | near-gene-3 | 654,589 | 0.00 | 3.16 | 3.16 | 0.55 | 0.66 | 0.41 |
|  | near-gene-5 | 2,487,192 | 0.00 | 3.16 | 3.16 | 0.54 | 0.65 | 0.41 |
|  | nonsense | 66,275 | 0.00 | 3.16 | 3.16 | 0.55 | 0.65 | 0.37 |
|  | splice-3 | 25,401 | 0.01 | 2.07 | 2.05 | 0.57 | 0.61 | 0.31 |
|  | splice-5 | 28,983 | 0.00 | 2.74 | 2.74 | 0.57 | 0.65 | 0.31 |
|  | stop-loss | 2,225 | 0.03 | 3.16 | 3.13 | 0.61 | 0.71 | 0.42 |
|  | unknown | 99,004,130 | 0.00 | 3.16 | 3.16 | 0.57 | 0.69 | 0.43 |
|  | untranslated-3 | 1,299,685 | 0.00 | 3.16 | 3.16 | 0.55 | 0.67 | 0.41 |
|  | untranslated-5 | 181,208 | 0.00 | 3.16 | 3.16 | 0.50 | 0.58 | 0.33 |

### Fine-mapping SLE-associated genomic regions using ΔMGW prioritization identifies potentially functional SNPs

To-date, more than 100 genomic loci have been associated with SLE.(43, 62) Here, we selected the genomic regions containing *FAM167A-BLK*, *STAT4*, and *TNIP1* for fine-mapping because these regions showed robust single-SNP associations (p < 5×10^−8^) with SLE in at least two ancestries (*FAM167A-BLK*: EA and AA; *STAT4*: EA and HA; *TNIP1*: EA and HA) and the association signals are not refined to a single SNP, due in part to strong linkage disequilibrium. Furthermore, neither the SNPs nor their LD proxies are protein-coding variants, leaving DNA topology as a potential functional mechanism. For each region, we first describe the previous SNP association results (43) and their LD patterns, by ancestry. Each region is then summarized by its ΔMGW measures which were used in frequentist and Bayesian ΔMGW-weighted analyses. SNPs identified by the ΔMGW-weighted analyses were subsequently investigated for existing functional evidence (See Methods and Materials).

### FAM167A-BLK

The SLE-associated region at 8p23 lies upstream of *FAM167A* and *BLK*, which are in a head-to-head gene orientation. Across the 500kb candidate region, 835 and 933 genotyped SNPs passed quality control in the EA and AA data, respectively. In the previous(43) logistic regression analyses, the primary peak of association was captured by a 60 kb window. In EA, the most significant SNP associations mapped to a 26 kb region of 16 SNPs in high LD (r^2^>0.8); within the AA data, the top associations were refined to a smaller 14 kb window containing 7 highly correlated SNPs (**Figure 5**).The summary statistics for ΔMGW for SNPs in the 500 kb and 60 kb regions were comparable to what was observed across the genome, with only a few SNPs imposing large changes in MGW (**Table S3**).

**Figure 5:**
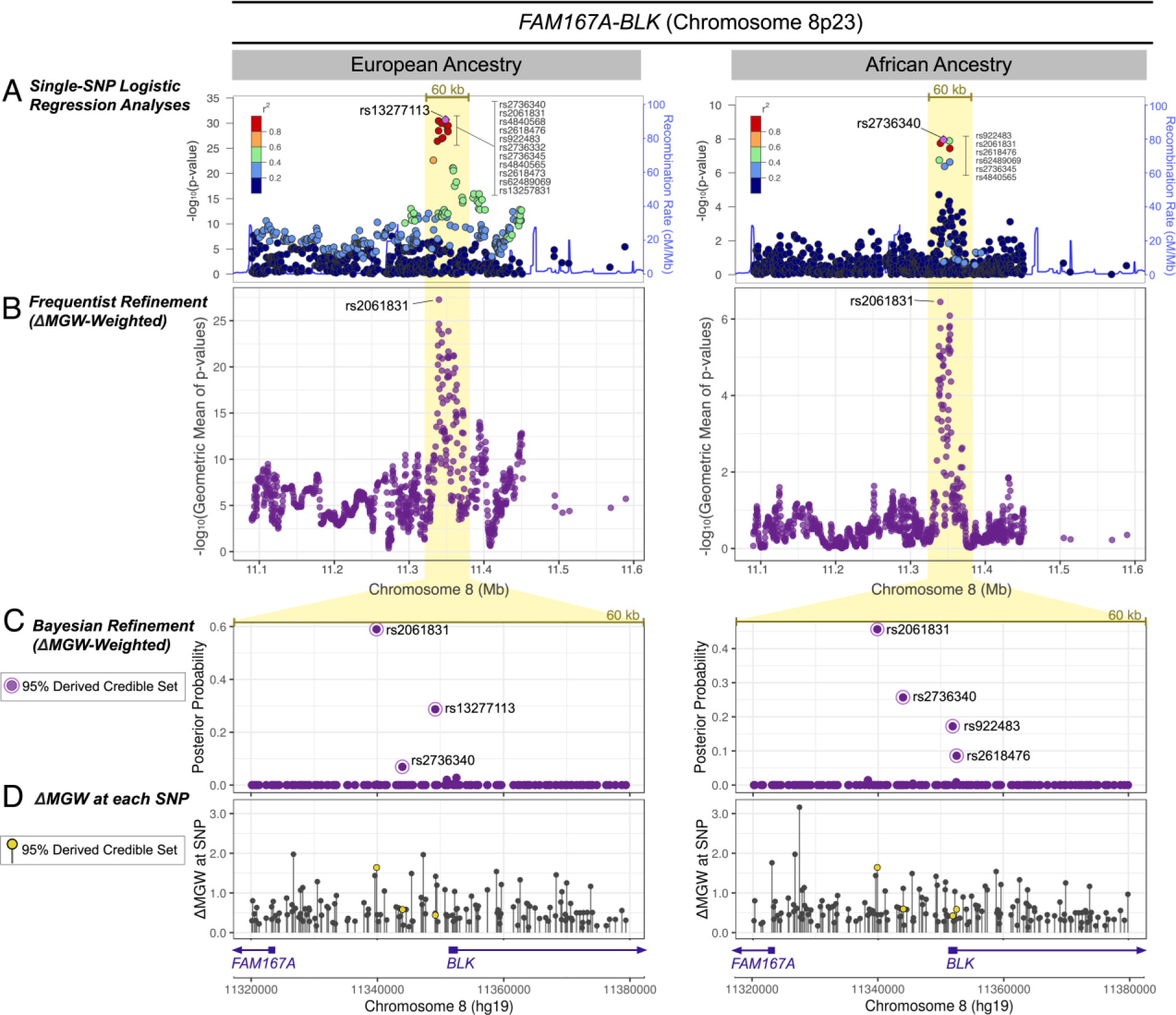
*FAM167A-BLK* ΔMGW prioritization by Frequentist and Bayesian Methods in European and African Ancestries. (A) Genotyped SNPs that passed quality control and were within 250kb of the top single-SNP association analysis in EA and AA data. A 60 kb region capturing the primary peak of association is highlighted. In both the EA and AA data a cluster of SNPs in high LD yielded the top association signals. (B) Using SKAT as a ΔMGW-weighted frequentist approach, rs2061831 was sharply prioritized over SNPs in the previously identified LD blocks. While the single-SNP logistic regression analyses in (A) identified a different top SNP in the EA (rs13277113) and AA (rs2736340) data, rs2061831 was consistently prioritized as the top SNP in both the EA and AA analyses. ΔMGW-weighting did not yield spurious associations for with SNPs outside the broad 60 kb peak of association highlighted in yellow. (C) SNPs within the 60 kb association peak were analyze by a Bayesian approach. The ΔMGW-weighted posterior probabilities are plotted. While the majority of SNPs yielded infinitesimal posterior probabilities, those comprising the 95% derived credible sets are labeled. Akin to the ΔMGW-weighted SKAT analyses, rs2061831 was again prioritized in both the EA and the AA data, with the largest posterior probability. (D) The ΔMGW is plotted for each SNP in the 60 kb region. The ΔMGW for a SNP is sequence-specific thus yielding the same values in EA and AA data. Differences between the two plots result from differences in genotyped SNP lists (i.e. SNPs that are monomorphic in one population would not be plotted). SNPs identified by the derived ΔMGW-weighted credible set are plotted in yellow. While rs2061831 had a large ΔMGW, other SNPs in the region had larger magnitudes of ΔMGW but did not show evidence of SLE-association. This illustrates the 2-parameter hypothesis of considering a combination of association signal and magnitude of ΔMGW. Prioritized SNPs fall upstream of both *FAM167A* and *BLK*.

Hypothesizing that plausibly functional SNPs can be identified by incorporating both ΔMGW and evidence for disease association, we applied two ΔMGW-weighted approaches via SKAT and Bayesian credible sets. For the 500 kb region, SKAT was applied in a 5-SNP rolling window (see Methods and Materials). Across the region, SNPs with the highest SKAT-weighted prioritizations largely followed the pattern observed in the single-SNP logistic regression analyses. That is, SNPs that were not previously associated with SLE were not prioritized solely on ΔMGW, as illustrated in the region outside of the 40 kb peak of association (**Figure 5**). When weighted by ΔMGW, rs2061831 was sharply prioritized in both the EA and AA analyses (**Figure 5**). In EA, rs2061831 was one of the 14 highly correlated SNPs identified by the single-SNP logistic regression analyses; likewise, in AA, it was also within the LD block comprising the 7 most highly associated SNPs. While the other SNPs in these LD blocks exhibited comparable SLE-association, rs2061831 had the greatest ΔMGW at 1.63 Å, prioritizing it above other SNPs in the weighted analyses. Importantly, while the single-SNP logistic regression analyses identified a different top SNP in EA (rs13277113) and AA (rs2736440), ΔMGW-weighting prioritized the same SNP (rs2061831), across ancestries. An unweighted SKAT prioritized the signal downstream of rs2061831, to the region where multiple SNPs from the same highly-associated LD block were included in the same 5-SNP windows (**Figure S1, Tables S4-S5**).

The ΔMGW-weighted frequentist fine-mapping evidence for rs2061831 was corroborated using the Bayesian refinement approach. In both EA and AA, the derived ΔMGW-weighted credible set placed the highest posterior probability on rs2061831 (58.9%-EA; 44.2%-AA) (**Figure 5**). In the un-weighted (standard) Bayesian analysis, rs2061831 was included in the EA (30.6% posterior probability) and AA (20.9% posterior probability) 95% credible sets, but it was not the highest prioritized **(Table S4-S5)**. Instead, the SNPs originally identified in the ancestry-specific logistic regression analyses were given the highest posterior probability—EA: rs13277113 (49.9% posterior probability), AA: rs2736340 (33.1%). Thus, like the frequentist approach, weighting by ΔMGW resolved the signal in both EA and AA to the same SNP, rs2061831.

Using ΔMGW as a prioritization metric, rs2061831 was consistently prioritized in both EA and AA data. SNP rs2061831 has a ΔMGW of 1.63 Å, which is 2 standard deviations above the mean across dbSNP150. Interestingly, this SNP is a transition polymorphism (Purine/Purine), a polymorphism type which we previously showed to have the smallest (on average) ΔMGW (**Table 1, Figure 3**). Considering only transition SNPs, rs2061831 is actually 4.52 standard deviations above the mean ΔMGW_transition SNPs_ (0.50 Å), indicating a considerable departure from the expected value and thus we would hypothesize a greater likelihood of functional relevance. Given the consistent evidence for a signal at rs2061831 in both the EA and AA data, we explored previously described (see Methods and Materials) functional data resources for evidence of biological relevance, in comparison to the top SNP signals from the single-SNP analyses (rs13277113 in EA; and rs2736440 in AA). All three SNPs are in high LD (R^2^>0.95) with one another in both EUR and AFR 1000 genomes data. Thus, it is unsurprising that all three SNPs yielded similar eQTL results via GTEx (data not shown). Despite the high LD, these three SNPs are physically separated by several kilobases. Of these three SNPs, rs2061831 is the only SNP that maps (via SCREEN) to a Candidate Cis-Regulatory Element (accession number: EH37E0941109) showing evidence for DNase, H3K27ac, and CTCF-binding activity. Consulting the 3D-genome browser yielded a larger number of long-range chromatin interactions in monocytes, B-Cells, and CD4 cells for rs2061831, in comparison to rs13277113 and rs2736440 (**Figure S2**). Thus, in this region, ΔMGW-weighting successfully differentiated among highly-correlated SNPs and prioritized rs2061831, a SNP within a potentially important regulatory region as documented by independent data.

### STAT4

The single-SNP SLE associations at 2q32 span the *STAT4* gene **(Figure 6)**. SNP associations reached genome significance in the EA and HA cohorts, with the strongest signals within intronic regions.(43) In the 500 kb region, there were 192 and 202 genotyped SNPs that passed quality control measures in EA and HA, respectively. In both ancestries, the primary peak of association was captured by a broad 110 kb window (**Figure 6**). The strongest associations in the EA data (p-values < 1×10^−62^) mapped to six SNPs in high LD, spanning 29 kb. Five of these SNPs also comprised the LD block of strongest associations in the HA data (p< 1×10^−13^), in a slightly narrower 26 kb region. The consistency of SNP association results in the EA and HA data provided a prime opportunity to test ΔMGW-prioritization among highly-correlated SNPs.

**Figure 6:**
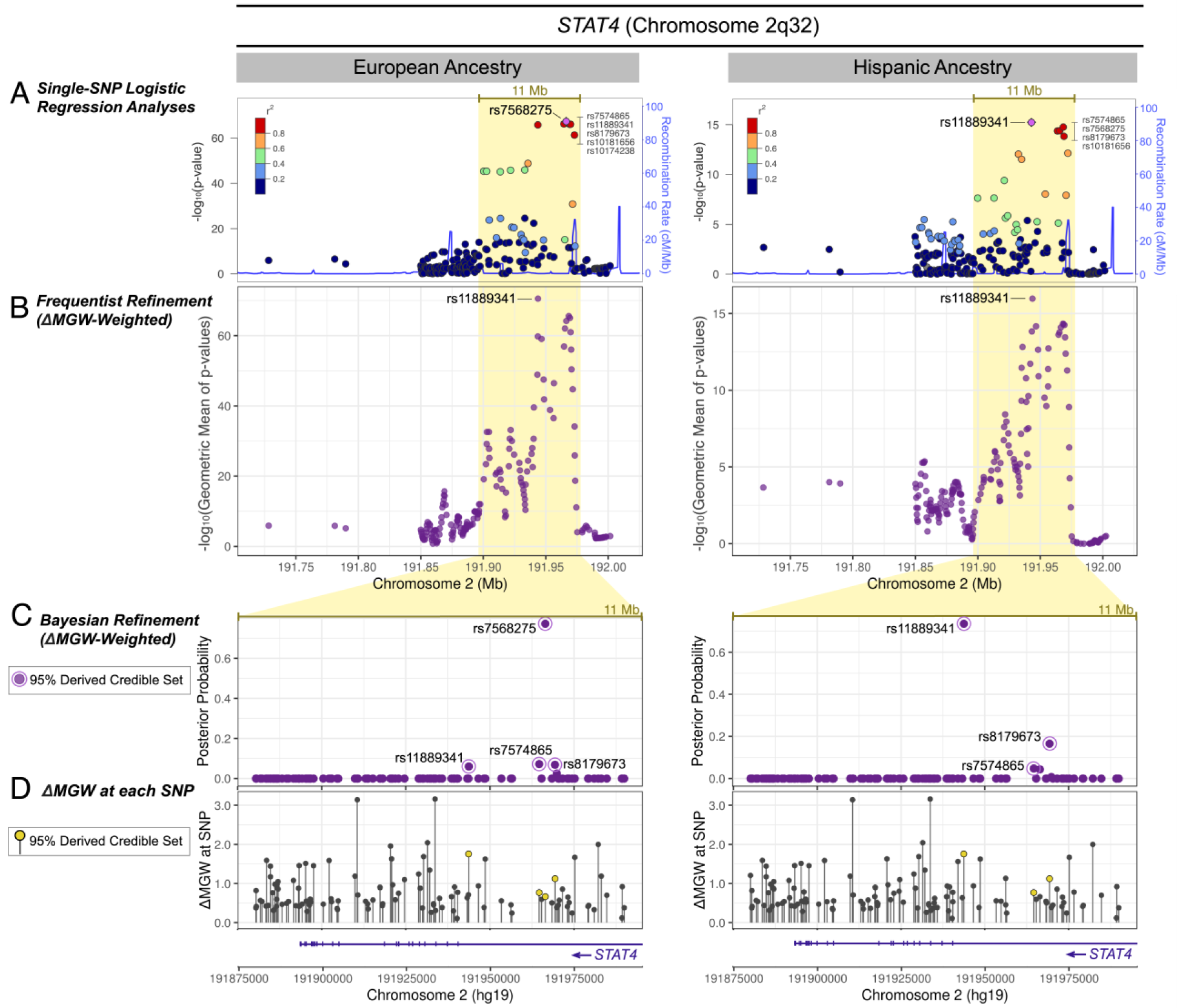
STAT4 ΔMGW prioritization by Frequentist and Bayesian Methods in European and Hispanic Ancestries. (A) Regional association plots in EA and HA for genotyped SNPs that passed quality control and were within 250kb of the top single-SNP association analysis in *STAT4.* Within the broad 11 Mb peak of association (highlighted in yellow), a cluster of SNPs in high LD yielded the top association values. (B) SNP refinement using SKAT with a ΔMGW-weighted approach sharply prioritizes rs11889341 in both EA and HA data. In the EA data, the ΔMGW-weighting shifted the top signal to rs1188931, whereas in the HA data, it simply further accentuated the signal above other SNPs. (C) For the highlighted 11 Mb region, SNP posterior probabilities are plotted for the derived, ΔMGW-weighted Bayesian analysis. While the frequentist MGW-weighted approach prioritized the same SNP (rs1188931) in both ancestries, this was not observed in the Bayesian approach. In the EA data, the Bayes factor for rs7568275 (BF=2.20×10^64^) was at such a large magnitude, that it was largely unaffected by ΔMGW-weighting. However, rs1188931 still entered the 95% derived credible set, but with a much smaller posterior probability (6.03%) compared to rs7568275 (77.25%). In the HA data, ΔMGW-weighting further prioritized rs1188931. (D) The ΔMGW for SNPs within the 11 Mb region. SNPs that were identified by the derived ΔMGW-weighted credible set are plotted in yellow. Again, the analytic approaches consider SNPs in the context of a 2-parameter hypothesis, evaluating SNPs for a combination of association signal and magnitude of ΔMGW. Hence, the prioritized SNPs (yellow) are not necessarily the SNPs with the largest ΔMGW in the region. Prioritized SNPs occur within an intron of *STAT4*.

The mean ΔMGW for SNPs in this region was 0.72 Å in EA and 0.73 Å in HA and both cohorts had a median ΔMGW of 0.56 Å. While these average ΔMGW were slightly higher than what was observed across the entire bi-allelic dbSNP dataset (mean=0.68 Å), the EA and HA medians were of the same magnitude (dbSNP ΔMGW median=0.56). The ΔMGW for SNPs within the 110 kb association window exhibited similar means as the 500 kb region (**Table S6**).

We again applied the two ΔMGW-weighted approaches using SKAT and Bayesian credible sets in the region. In EA, the ΔMGW-weighted SKAT analyses shifted the top signal upstream to rs11889341, which markedly increased its priority (**Figure 6**). This SNP was one of the top six SNPs in the single-SNP association LD-block. While it and the other five SNPs were all significantly associated with SLE, rs11889341 had the greatest ΔMGW at 1.75 Å, which prioritized it over the other SNPs in the LD block; the remaining SNPs had ΔMGW values ranging from 0.31-1.12 Å (**Figure 6**). In HA, weighting by ΔMGW in the SKAT analysis also prioritized rs11889341 as the top SNP. This SNP was previously identified with the best p-value in the single-SNP association analysis, but in the ΔMGW-weighted approach, its prioritization distinctly increased relative to the other SNPs in the LD block (**Figure 6**).

In the Bayesian analysis, rs11889341 was included in the EA and HA derived ΔMGW-weighted 95% credible sets (**Figure 6**). In EA, rs11889341 was not in the unweighted 95% credible set but inclusion of ΔMGW increased its posterior probability from 2.4% to 6.0% (**Table S7, Figure S3**). In EA, rs7568275 yielded the strongest signal in both the unweighted (81.0% posterior probability) and derived ΔMGW-weighted (77.3% posterior probability) credible sets (**Table S7**). This is important to note, as rs7568275 had a much smaller ΔMGW (0.66 Å) than rs11889341 (1.75 Å.). This provided an example where the magnitude of the Bayes factor was so large (p=4×10^68^), that the influence of ΔMGW was largely diminished in the analysis. However, despite the predominant rs7568275 signal, the derived credible set still detected rs11889341, the SNP identified by the ΔMGW-weighted SKAT approach. In the HA data, rs11889341 yielded the largest posterior probability in the ΔMGW-weighted derived credible set. This SNP also had the largest posterior probability in the unweighted credible set. Unlike the EA analysis, where the magnitude of the Bayes factor dominated the impact of the ΔMGW-weighting, in the HA data, the ΔMGW strongly increased the posterior probability of rs11889341 from 58.6% to 73.5% (**Figure 6, Table S8**). This limited the derived 95% credible set to only 3 SNPs: rs11889341 (73.5%), rs8179673 (16.6%), and rs7574865 (4.8%) (**Table S8**).

In the single-SNP association analyses of *STAT4* SNPs, the association signal was refined to an LD block of 6 SNPs in the EA data and 5 SNPs in the HA dataset. In ΔMGW-weighted analyses, rs11889341 was sharply prioritized over other SNPs in the LD block, with an exception in the EA ΔMGW-weighted derived credible set, where the high magnitude of the Bayes factor for rs7568275 (bf=2.20×10^64^) over other SNPs (bf <=1.79×10^63^) largely negated any impact of ΔMGW in this analysis. Considering the evidence for rs11889341 in the other three analyses due to its strong combination of SLE association and ΔMGW, we would hypothesize that rs11889341 would be a candidate functional polymorphism. Like rs2061831 in *FAM167A-BLK*, rs11889341 is also a transition SNP (purine/purine). While transition SNPs are more frequent across the genome (previously shown in Table S1), there are few transition SNPs (+/− 4 nucleotides) that yield such a high ΔMGW (mean ΔMGW for transition SNPs=0.50 Å). Evaluation of publically available functional datasets (see METHODS) yielded limited information for both rs7568275 and rs11889341. Neither of these SNPs were identified as eQTLs in GTEx nor were they within Candidate Cis-Regulatory regions (cCREs). Furthermore, neither variant was shown with long range chromatin interactions in the in the currently available HI-C data via the 3D genome browser. However, despite the lack of functional information from these resources, functional evaluation of rs11889341 is available via a 2018 study by Patel and colleagues, where transancestral mapping identified rs11889341 with strong association with SLE.(63) In this study, rs11889341 was associated with *STAT1* expression in B-cells through increased binding of the transcription factor, HMGA1. Given the relationship between transcription factor binding and DNA topology(20, 31, 32, 64, 65), we hypothesize that the identified functional activity of rs11889341 (via HMGA1 binding) may be mediated by the large MGW change imposed by the SNP’s alleles.

### TNIP1

Previous single-SNP association analyses(43) identified genome-wide significant findings (p<5×10^−8^) in EA and HA data at 5q33 (**Figure 7**). In the 500 kb region, there were 497 and 500 high quality genotyped SNPs in the EA and HA data, respectively. The peak of SLE association is captured by a 40 kb window which encompasses most of the *TNIP1* gene. In the EA data, the top associations mapped to three SNPs (rs960709, rs10036748, rs6889239) in high LD, spanning 3 kb of a *TNIP1* intron. These three SNPs are also encompassed by the associated LD block in the HA data, where four, highly correlated SNPs (rs1422673, rs960709, rs10036748, and rs6889239) yielded p-values < 5×10^−8^. As completed in the *FAM167A-BLK* and *STAT4* regions, we again applied ΔMGW-weighted fine-mapping strategies to prioritize these non-coding SLE-associated SNPs.

**Figure 7:**
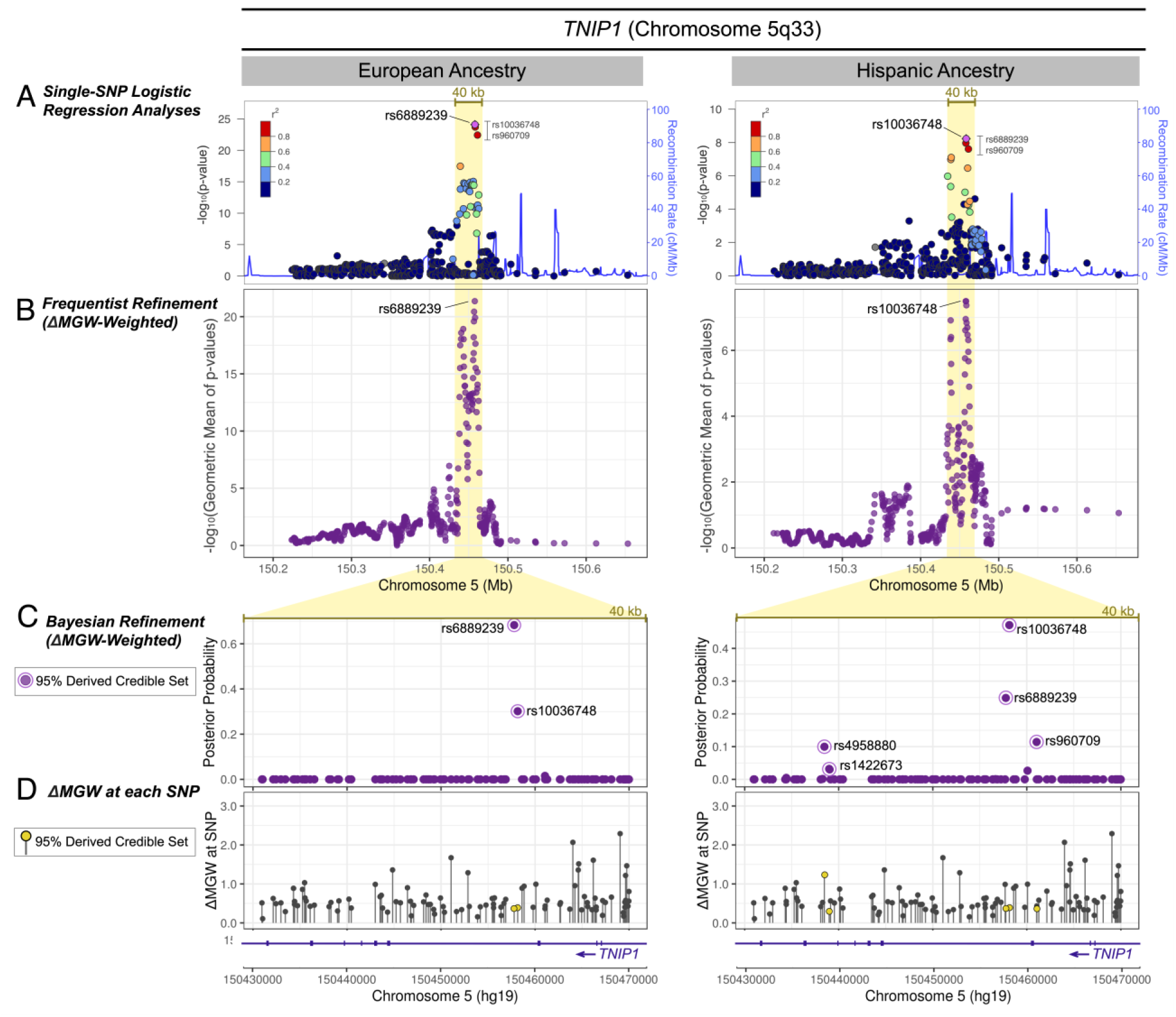
*TNIP*1 ΔMGW prioritization by Frequentist and Bayesian Methods in European and Hispanic Ancestries. (A) Genotyped SNPs within 250 kb of the top single-SNP association analysis are shown for EA and HA. The 40 kb region that captures the primary peak of association is highlighted in yellow. In EA and HA, the same three SNPs (rs10036748, rs6889239, and rs960709) show the highest association values and are all in high LD. In EA rs6889239 has the best p-value and rs10036748 yields the best p-value in HA. (B) Analyzing the region with SKAT in a ΔMGW-weighted approach. In this region, for these SNPs, including ΔMGW did not provide differential prioritization, rs6889239 remained the top signal for EA and rs10036748 for HA. (C) For each SNP in the 40 kb region, the posterior probabilities are plotted for the derived, ΔMGW-weighted Bayesian analysis. The weighted Bayesian analysis did not alter the relative signals observed in the single-SNP logistic regression analyses. In the EA data, rs6889239 yielded the largest posterior probability in EA and rs10036748 remained the top signal for HA. (D) The ΔMGW is plotted for each genotyped SNP that passed quality control measures. SNPs that were identified by the derived ΔMGW-weighted credible set are plotted in yellow. These prioritized SNPs have comparatively low magnitudes of ΔMGW, indicating that the driving factor of these SNP prioritizations stemmed from their SLE associations and not their magnitude of ΔMGW.

In the *TNIP3* region, the lists of high-quality genotyped SNPs were largely the same between the EA and HA datasets. Consequently, the statistics for ΔMGW in this region were very similar between the two cohorts. Across the 500 kb window of high quality SNPs, the average ΔMGW was 0.67 Å (median=0.55 Å) in both EA and HA. (**Table S9**) These values were slightly lower than the observed mean for bi-allelic SNPs from dbSNP (**Table 1**).

The SKAT analyses yielded similar results between the EA and HA data. The ΔMGW-weighted analyses did not effectively prioritize or refine the SNP signal. Unlike *FAM167A-BLK* and *STAT4*, ΔMGW-weighting did not resolve the top signal to the same SNP in both ancestries. Instead, in *TNIP1*, the top SNPs in the ΔMGW-weighted analyses for EA (rs6889239) and HA (rs10036748) were the same as those identified in the single-SNP logistic regression analysis (**Figure 7**). The SNPs that were prioritized in the unweighted SKAT analyses were also prioritized in the ΔMGW-weighted analyses; notably, in this region ΔMGW-weighting actually dampened the signal because the SNPs with the greatest SLE association values had low magnitudes of ΔMGW (ranging from 0.31-0.37 Å). This pattern was also observed in the Bayesian approach, where SNPs with the highest posterior probabilities in the derived credible sets exhibited lower posterior probabilities than in the unweighted credible set **(Figures 7 and S4 and Tables S10-S11**), again due to the low magnitudes of ΔMGW for top-associated SNPs.

In *TNIP1*, the ΔMGW-weighted analyses did not differentially prioritize SNPs in comparison to the unweighted approaches. While there were SNPs with large ΔMGW in the region, these did not have strong SLE-associations. Unlike the *FAM167A-BLK* and *STAT4* regions, where ΔMGW successfully prioritized specific SNPs, this was not achieved in the *TNIP1* region. This could indicate several possibilities, including: ΔMGW may not be a relevant mechanism for these SNPs, another DNA measure may be more informative, DNA topology may not be a functional driver for this region, and/or or the functional variant was not included in these analyses. Here, an alternative strategy is required to identify the most plausible functional polymorphisms.

## Discussion

Sequence-dependent DNA topology could provide important functional context for associations, especially for polymorphisms that do not impose protein changes (e.g., coding-synonymous) and/or variants mapping to non-coding regions. We explored ΔMGW, a specific sequence-dependent measure of DNA topology, as a weighting variable in fine-mapping analyses. In a sample of 300k SNPs, Wang *et al*. previously found that MGW-preserving SNPs are more common.(42) Here, we built upon these findings through a full census of bi-allelic SNPs (n=199,038,197) across the genome. We showed the observed genomic ΔMGW was significantly lower than the complete ΔMGW sample space. These findings were consistent with the relative frequencies of transversion (∼33%) and transition (∼66%) mutations in the human genome.(59, 60) We hypothesized that phenotypically-associated SNPs with large ΔMGW would be more likely to impose functional consequences; and thus, proposed ΔMGW as a prioritization metric in fine-mapping studies.

We tested our hypothesis using ΔMGW weights in two fine-mapping approaches in three regions (*FAM167A-BLK*, *STAT4*, and *TNIP1*) with well-established SLE associations. In *FAM167A-BLK and STAT4*, we successfully identified SNPs of possible functional consequence, underscoring ΔMGW as a plausibly informative prioritization metric in fine-mapping studies.

There are several advantages to using sequence dependent topology, such as ΔMGW, as a weighting metric in fine-mapping studies. For one, it is an intrinsic variable, inherent to the genetic sequence surrounding the polymorphism; thus, it is not reliant on external data which may offer limited information for the SNPs of interest (database bias). As an intrinsic variable it is also not ancestry specific, tissue specific, or sample size dependent. Limitations in external (non-intrinsic) data may down-weight potentially causal SNPs due to a lack of available functional data. While publically available functional resources continue to expand, they still present these challenges, especially for rare or novel variants. This is particularly relevant for diverse study populations where annotation resources based on European data offer inadequate or no coverage for regions of interest.(14) For example, Sherman *et al*. presented deep sequencing in 910 individuals of African descent and found over 296 million base pairs which were absent in the human reference genome.(15) Novel variants or regions are unlikely to be annotated by commonly used resources. Therefore, while a SNP’s functional relevance can be supported by public resources, a lack of information does not necessarily indicate a variant’s lack of function. This was illustrated by rs11889341 in *STAT4*, which lacked functional information from public resources (GTEx, ENCODE, 3D-genome browser)(55, 56, 58), but in a targeted functional study by Patel *et al*., rs11889341 was correlated with gene expression and binding of the transcription factor HMGA1.(63) We identified rs11889341 using ΔMGW as the prioritizing variable. Thus, prioritizing SNPs by a factor intrinsic to DNA may help alleviate some bias that would otherwise be introduced by missing data from publically available functional datasets. Consequently, we propose including ΔMGW among annotation resources used in SNP-weighted fine-mapping methods.

Changes in DNA topology can potentially impact an array of biological functions such as transcription factor binding, chromatin remodeling, or methylation.(20, 21, 23, 26, 31, 32, 36) Likewise, using DNA topology as a SNP prioritization metric does not limit functional information to a single biological mechanism. This may be especially beneficial when the relationship between phenotype and biological mechanism is unknown. While functional work in *STAT4* showed that rs11889341 altered HMGA1 binding, functional work is still needed to evaluate the rs2061831 genotype in *FAM167A-BLK*. Here, the biological implications of rs2061831 could involve transcription factor binding, and/or, given its apparent location within a long-range chromatin interaction hotspot (**Figure S1**), chromatin organization. Considering the strong trans-ancestral signal of rs2061831 across EA and AA, further functional work should explore whether this SNP acts through an independent functional mechanism or through interactions with other variants in the region (e.g. within the context of sequence-dependent structural motifs), such as the insertion-deletion identified in a study of ATAC-seq data in 100 individuals of British Ancestry.(66) Leveraging changes in DNA topology can identify potentially causal polymorphisms and also generate specific hypotheses for functional follow-up studies. Furthermore, sequence-dependent DNA topology is a weighting scheme that informatively decouples SNPs in high LD, a long sought after feature as associations and eQTLs are often confounded by LD. In *FAM167A-BLK*, we observed comparable eQTL evidence for SNPs in the associated LD cluster, making eQTL status ineffective at differentiating highly-correlated SNPs. Instead, consideration of sequence-dependent ΔMGW allowed differential prioritization among these otherwise, highly-correlated SNPS, selecting rs2061831 as a plausible functional candidate SNP.

Another advantage to using local DNA topology in fine-mapping studies is its consistency of information across ancestries. Assuming identical flanking sequence (e.g., no genomic variant within +/− 4 bases of the SNP), a SNP’s impact on DNA topology would be constant across ancestries, highlighting the potential utility of DNA topology as a means of resolving association signals across ancestries. Here, we showed that ΔMGW-weighted analyses of *FAM167A*-*BLK* and *STAT4* resolved the association signal to the same SNP in each ancestry via the frequentist approach, followed by largely corroborating evidence via the derived credible sets in the Bayesian approach. Notably, rs2061831 was not the top-associated SNP in either the ancestry-specific analyses; however, it was previously identified via the SLE Immunochip trans-ancestral meta-analysis, where combining association signals across ancestries identified it as the top SNP.(43)

### Limitations and Future Work

There are several considerations and limitations to using sequence-dependent topology as a weighting metric in fine-mapping analyses. Notably, some of these limitations could result in inconclusive and/or insignificant results, as observed in the *TNIP1* region. First, the functional variants may not have been genotyped or imputed in the study. Analyses that utilize SNP-specific weights decouple associations from LD. Thus, a weighted metric performs best when the functional SNP is included in the analysis set. For this reason, we propose application of this prioritization technique in genomic regions where there is high confidence that the functional variants have been genotyped or imputed. We note this limitation exists for any statistical association method.

Second, DNA topology, here ΔMGW, may not be the mechanism impacting phenotype. While sequence dependent DNA topology can influence a number of functional factors(18, 21, 23, 24, 32), it is not the only source of biological interactions and could be irrelevant for a specific phenotype. Thus, when using change in DNA topology, such as ΔMGW, in fine-mapping studies, analyses should be considered in the form of a two-parameter hypothesis – a combination of association signal and ΔMGW. For example, in both the *FAM167A-BLK* and *STAT4* regions, the highest prioritized SNPs, rs2061831 and rs11889341, did not have the largest magnitude of ΔMGW in the regions (**Figures 5-6**). Instead, these two SNPs were prioritized by their combined SLE-association and ΔMGW.

Third, we placed greater weights on SNPs with larger magnitudes of change on DNA topology. We recognize that even small changes could yield functional consequences. Thus, future studies should explore weighting SNPs by particular topological profiles (e.g., those matching binding site profiles). For instance, our *TNIP1* analyses did not show strong signals when weighting by the magnitude of ΔMGW, but this does not definitively rule out MGW as a functional mechanism (e.g. driven by pattern, not magnitude). The focus on MGW was motivated by the breadth of study on MGW and function.(18, 20, 32, 34, 36) So while this manuscript considered a single parameter, ΔMGW, we are currently expanding to incorporate additional measures (e.g., helix twist, roll) through multivariate approaches that account for the correlation structure (dependencies) among spatial measures.

Fourth, in this study, we used SKAT and a derived credible sets (Bayesian) approach to apply a topological weighting scheme to prioritize SNPs; however, we note that there are other methods that can incorporate weights for SNP association analyses.(10, 67) Here, we assumed that the majority of variants in the region are non-causal, which is why we selected SKAT over a combined burden test. However, we note that the results from SKAT and SKAT-O were largely similar. Similarly, in case of the Bayesian approach applied here, a limitation is its assumption that a single causal SNP exists in a region, but other Bayesian methods can be explored.(53, 68) In the EA *STAT4* data, the magnitudes of the Bayes factors were so large that weighting by ΔMGW yielded minimal impact. Future work should consider approaches to scale weighting schemes by a constant derived from the magnitude of signal across a genomic region. In the SKAT approach, for the sliding analysis window, we used five SNPs, which should yield a region that is neither too wide nor too unstable. Additional testing could potentially improve optimization of parameters for this analysis. Furthermore, we emphasize that our evaluation of the SKAT results by summarizing each SNP as the geometric mean of SKAT-analysis p-values should be regarded as a metric for prioritizing SNPs, not an association analyses, as these values do not have the statistical properties of a p-value. Overall, these limitations should be carefully considered when applying these specific methods; but they also highlight opportunities to further explore the relationship between sequence-dependent DNA topology and phenotype associations.

In summary, weighting SNP associations by functional data can greatly improve identification of potentially causal SNPs; however, existing annotation resources can negatively affect these outcomes when SNP information is unavailable in public datasets, especially in non-EA populations.(8, 10, 11, 14) In this study, we presented and tested sequence-dependent DNA topology as a novel annotation source for genetic fine-mapping studies. As an intrinsic property, sequence-dependent DNA shape alleviates many of the challenges imposed by external data resources; and it provides potential functional (testable) context for associations (e.g. topological disruption for protein binding). Using ΔMGW in weighted analyses, we successfully prioritized functional SNPs in two SLE-associated regions with high LD. Likewise, as an annotation resource, sequence-dependent DNA topology, such as ΔMGW, is readily applicable in any fine-mapping methods that can incorporate continuous values for SNP weights. Altogether, this manuscript presents methods that are immediately applicable to existing genetic data, and it illustrates how sequence-dependent DNA topology can be used as a paradigm to investigate and understand genetic associations in fine-mapping studies.

## Funding

This work was supported by the National Institutes of Health [HG007112-01, U01 NS036695]; and the National Aeronautics and Space Administration [NNX16A069A].

## Declaration of Interests

The authors declare no competing interests.

## Acknowledgements

We thank M.A. Espeland, M.A. Alexander-Miller, B.I. Freedman, K.D. Zimmerman, M.C. Marion, and M.E. Comeau for discussions on content and feedback on the work in this paper. HCA was also supported through the Wake Forest Biomedical Sciences Graduate School.

## Notes

#### Summary of Updates

Figure one was changed to better convey concept of sequence-dependent DNA shape. Minor formatting edits were made to Figures 2 and 3. A Graphical abstract has been added. Some minor clarifying points have been included in the introduction and discussion.

## References

1. MacArthur, J., Bowler, E., Cerezo, M., Gil, L., Hall, P., Hastings, E., Junkins, H., McMahon, A., Milano, A., Morales, J., et al. (2017) The new NHGRI-EBI Catalog of published genome-wide association studies (GWAS Catalog). Nucleic Acids Res., 45, D896–D901.

2. Visscher, P.M., Brown, M.A., McCarthy, M.I. and Yang, J. (2012) Five years of GWAS discovery. Am. J. Hum. Genet., 90, 7–24.

3. Visscher, P.M., Wray, N.R., Zhang, Q., Sklar, P., McCarthy, M.I., Brown, M.A. and Yang, J. (2017) 10 Years of GWAS Discovery: Biology, Function, and Translation. Am. J. Hum. Genet., 101, 5–22.

4. McCarthy, M.I., Abecasis, G.R., Cardon, L.R., Goldstein, D.B., Little, J., Ioannidis, J.P.A. and Hirschhorn, J.N. (2008) Genome-wide association studies for complex traits: consensus, uncertainty and challenges. Nat. Rev. Genet., 9, 356–369.

5. Manolio, T.A., Collins, F.S., Cox, N.J., Goldstein, D.B., Hindorff, L.A., Hunter, D.J., McCarthy, M.I., Ramos, E.M., Cardon, L.R., Chakravarti, A., et al. (2009) Finding the missing heritability of complex diseases. Nature, 461, 747–753.

6. Pasaniuc, B. and Price, A.L. (2017) Dissecting the genetics of complex traits using summary association statistics. Nat. Rev. Genet., 18, 117–127.

7. Farh, K.K.-H., Marson, A., Zhu, J., Kleinewietfeld, M., Housley, W.J., Beik, S., Shoresh, N., Whitton, H., Ryan, R.J.H., Shishkin, A.A., et al. (2015) Genetic and epigenetic fine mapping of causal autoimmune disease variants. Nature, 518, 337–343.

8. Gomez-Cabrero, D., Abugessaisa, I., Maier, D., Teschendorff, A., Merkenschlager, M., Gisel, A., Ballestar, E., Bongcam-Rudloff, E., Conesa, A. and Tegnér, J. (2014) Data integration in the era of omics: current and future challenges. BMC Syst. Biol., 8, I1.

9. Faye, L.L., Machiela, M.J., Kraft, P., Bull, S.B. and Sun, L. (2013) Re-Ranking Sequencing Variants in the Post-GWAS Era for Accurate Causal Variant Identification. PLOS Genet., 9, e1003609.

10. Kichaev, G., Yang, W.-Y., Lindstrom, S., Hormozdiari, F., Eskin, E., Price, A.L., Kraft, P. and Pasaniuc, B. (2014) Integrating Functional Data to Prioritize Causal Variants in Statistical Fine-Mapping Studies. PLOS Genet., 10, e1004722.

11. Xu, Z. and Taylor, J.A. (2009) SNPinfo: integrating GWAS and candidate gene information into functional SNP selection for genetic association studies. Nucleic Acids Res., 37, W600–W605.

12. Lee, S., Wu, M.C. and Lin, X. (2012) Optimal tests for rare variant effects in sequencing association studies. Biostatistics, 13, 762–775.

13. Nicolae, D.L., Gamazon, E., Zhang, W., Duan, S., Dolan, M.E. and Cox, N.J. (2010) Trait-Associated SNPs Are More Likely to Be eQTLs: Annotation to Enhance Discovery from GWAS. PLOS Genet., 6, e1000888.

14. Kessler, M.D., Yerges-Armstrong, L., Taub, M.A., Shetty, A.C., Maloney, K., Jeng, L.J.B., Ruczinski, I., Levin, A.M., Williams, L.K., Beaty, T.H., et al. (2016) Challenges and disparities in the application of personalized genomic medicine to populations with African ancestry. Nat. Commun., 7, 12521.

15. Sherman, R.M., Forman, J., Antonescu, V., Puiu, D., Daya, M., Rafaels, N., Boorgula, M.P., Chavan, S., Vergara, C., Ortega, V.E., et al. (2019) Assembly of a pan-genome from deep sequencing of 910 humans of African descent. Nat. Genet., 51, 30–35.

16. Need, A.C. and Goldstein, D.B. (2009) Next generation disparities in human genomics: concerns and remedies. Trends Genet. TIG, 25, 489–494.

17. Manrai, A.K., Funke, B.H., Rehm, H.L., Olesen, M.S., Maron, B.A., Szolovits, P., Margulies, D.M., Loscalzo, J. and Kohane, I.S. (2016) Genetic Misdiagnoses and the Potential for Health Disparities. N. Engl. J. Med., 375, 655–665.

18. Privalov, P.L., Dragan, A.I., Crane-Robinson, C., Breslauer, K.J., Remeta, D.P. and Minetti, C.A.S.A. (2007) What Drives Proteins into the Major or Minor Grooves of DNA? J. Mol. Biol., 365, 1–9.

19. Yakovchuk, P., Protozanova, E. and Frank-Kamenetskii, M.D. (2006) Base-stacking and base-pairing contributions into thermal stability of the DNA double helix. Nucleic Acids Res., 34, 564–574.

20. Yang, L., Orenstein, Y., Jolma, A., Yin, Y., Taipale, J., Shamir, R. and Rohs, R. (2017) Transcription factor family-specific DNA shape readout revealed by quantitative specificity models. Mol. Syst. Biol., 13, 910.

21. Duan, C., Huan, Q., Chen, X., Wu, S., Carey, L.B., He, X. and Qian, W. (2018) Reduced intrinsic DNA curvature leads to increased mutation rate. Genome Biol., 19, 132.

22. Sati, S. and Cavalli, G. (2017) Chromosome conformation capture technologies and their impact in understanding genome function. Chromosoma, 126, 33–44.

23. Lazarovici, A., Zhou, T., Shafer, A., Dantas Machado, A.C., Riley, T.R., Sandstrom, R., Sabo, P.J., Lu, Y., Rohs, R., Stamatoyannopoulos, J.A., et al. (2013) Probing DNA shape and methylation state on a genomic scale with DNase I. Proc. Natl. Acad. Sci. U. S. A., 110, 6376–6381.

24. Abe, N., Dror, I., Yang, L., Slattery, M., Zhou, T., Bussemaker, H.J., Rohs, R. and Mann, R.S. (2015) Deconvolving the recognition of DNA shape from sequence. Cell, 161, 307–318.

25. Bansal, M., Kumar, A. and Yella, V.R. (2014) Role of DNA sequence based structural features of promoters in transcription initiation and gene expression. Curr. Opin. Struct. Biol., 25, 77–85.

26. Parker, S. and Tullius, T.D. (2011) DNA shape, genetic codes, and evolution. Curr. Opin. Struct. Biol.

27. Olson, W.K., Bansal, M., Burley, S.K., Dickerson, R.E., Gerstein, M., Harvey, S.C., Heinemann, U., Lu, X.-J., Neidle, S., Shakked, Z., et al. (2001) A Standard Reference Frame for the Description of Nucleic Acid Base-pair Geometry. J. Mol. Biol., 313, 229–237.

28. Lu, X.-J. and Olson, W.K. (1999) Resolving the discrepancies among nucleic acid conformational analyses11Edited by I. Tinoco. J. Mol. Biol., 285, 1563–1575.

29. Dickerson, R.E. (1989) Definitions and nomenclature of nucleic acid structure components. Nucleic Acids Res., 17, 1797–1803.

30. Rohs, R., West, S.M., Sosinsky, A., Liu, P., Mann, R.S. and Honig, B. (2009) The role of DNA shape in protein-DNA recognition. Nature, 461, 1248–1253.

31. Meysman, P., Marchal, K. and Engelen, K. (2012) DNA structural properties in the classification of genomic transcription regulation elements. Bioinforma. Biol. Insights, 6, 155–168.

32. Stella, S., Cascio, D. and Johnson, R.C. (2010) The shape of the DNA minor groove directs binding by the DNA-bending protein Fis. Genes Dev., 24, 814–826.

33. Irobalieva, R.N., Fogg, J.M., Catanese, D.J., Catanese, D.J., Sutthibutpong, T., Chen, M., Barker, A.K., Ludtke, S.J., Harris, S.A., Schmid, M.F., et al. (2015) Structural diversity of supercoiled DNA. Nat. Commun., 6, 8440.

34. Morgunova, E., Yin, Y., Jolma, A., Dave, K., Schmierer, B., Popov, A., Eremina, N., Nilsson, L. and Taipale, J. (2015) Structural insights into the DNA-binding specificity of E2F family transcription factors. Nat. Commun., 6, 10050.

35. Ngo, T.T.M., Zhang, Q., Zhou, R., Yodh, J.G. and Ha, T. (2015) Asymmetric unwrapping of nucleosomes under tension directed by DNA local flexibility. Cell, 160, 1135–1144.

36. Perino, M., van Mierlo, G., Karemaker, I.D., van Genesen, S., Vermeulen, M., Marks, H., van Heeringen, S.J. and Veenstra, G.J.C. (2018) MTF2 recruits Polycomb Repressive Complex 2 by helical-shape-selective DNA binding. Nat. Genet., 50, 1002–1010.

37. Chen, C. and Pettitt, B.M. (2016) DNA Shape versus Sequence Variations in the Protein Binding Process. Biophys. J., 110, 534–544.

38. Shepherd, J.W., Greenall, R.J., Probert, M.I.J., Noy, A. and Leake, M.C. (2020) The emergence of sequence-dependent structural motifs in stretched, torsionally constrained DNA. Nucleic Acids Res., 10.1093/nar/gkz1227.

39. Chiu, T.-P., Comoglio, F., Zhou, T., Yang, L., Paro, R. and Rohs, R. (2016) DNAshapeR: an R/Bioconductor package for DNA shape prediction and feature encoding. Bioinformatics, 32, 1211–1213.

40. Zhou, T., Shen, N., Yang, L., Abe, N., Horton, J., Mann, R.S., Bussemaker, H.J., Gordân, R. and Rohs, R. (2015) Quantitative modeling of transcription factor binding specificities using DNA shape. Proc. Natl. Acad. Sci., 112, 4654–4659.

41. Duzdevich, D., Redding, S. and Greene, E. (2014) DNA Dynamics and Single-Molecule Biology. Chem. Rev., 114, 3072–3086.

42. Wang, X., Zhou, T., Wunderlich, Z., Maurano, M.T., DePace, A.H., Nuzhdin, S.V. and Rohs, R. (2018) Analysis of Genetic Variation Indicates DNA Shape Involvement in Purifying Selection. Mol. Biol. Evol., 35, 1958–1967.

43. Langefeld, C.D., Ainsworth, H.C., Graham, D.S.C., Kelly, J.A., Comeau, M.E., Marion, M.C., Howard, T.D., Ramos, P.S., Croker, J.A., Morris, D.L., et al. (2017) Transancestral mapping and genetic load in systemic lupus erythematosus. Nat. Commun., 8, 16021.

44. van Dijk, M. and Bonvin, A.M.J.J. (2009) 3D-DART: a DNA structure modelling server. Nucleic Acids Res., 37, W235–239.

45. Pettersen, E.F., Goddard, T.D., Huang, C.C., Couch, G.S., Greenblatt, D.M., Meng, E.C. and Ferrin, T.E. (2004) UCSF Chimera--a visualization system for exploratory research and analysis. J. Comput. Chem., 25, 1605–1612.

46. Haeussler, M., Zweig, A.S., Tyner, C., Speir, M.L., Rosenbloom, K.R., Raney, B.J., Lee, C.M., Lee, B.T., Hinrichs, A.S., Gonzalez, J.N., et al. (2019) The UCSC Genome Browser database: 2019 update. Nucleic Acids Res., 47, D853–D858.

47. Karolchik, D., Hinrichs, A.S., Furey, T.S., Roskin, K.M., Sugnet, C.W., Haussler, D. and Kent, W.J. (2004) The UCSC Table Browser data retrieval tool. Nucleic Acids Res., 32, D493–D496.

48. Lander, E.S., Linton, L.M., Birren, B., Nusbaum, C., Zody, M.C., Baldwin, J., Devon, K., Dewar, K., Doyle, M., FitzHugh, W., et al. (2001) Initial sequencing and analysis of the human genome. Nature, 409, 860–921.

49. Wu, M.C., Lee, S., Cai, T., Li, Y., Boehnke, M. and Lin, X. (2011) Rare-variant association testing for sequencing data with the sequence kernel association test. Am. J. Hum. Genet., 89, 82–93.

50. Lee, S., Emond, M.J., Bamshad, M.J., Barnes, K.C., Rieder, M.J., Nickerson, D.A., NHLBI GO Exome Sequencing Project—ESP Lung Project Team, Christiani, D.C., Wurfel, M.M. and Lin, X. (2012) Optimal unified approach for rare-variant association testing with application to small-sample case-control whole-exome sequencing studies. Am. J. Hum. Genet., 91, 224–237.

51. Stephens, M. and Balding, D.J. (2009) Bayesian statistical methods for genetic association studies. Nat. Rev. Genet., 10, 681–690.

52. Marchini, J., Howie, B., Myers, S., McVean, G. and Donnelly, P. (2007) A new multipoint method for genome-wide association studies by imputation of genotypes. Nat. Genet., 39, 906–913.

53. The Wellcome Trust Case Control Consortium, Maller, J.B., McVean, G., Byrnes, J., Vukcevic, D., Palin, K., Su, Z., Howson, J.M.M., Auton, A., Myers, S., et al. (2012) Bayesian refinement of association signals for 14 loci in 3 common diseases. Nat. Genet., 44, 1294–1301.

54. Kichaev, G., Roytman, M., Johnson, R., Eskin, E., Lindström, S., Kraft, P. and Pasaniuc, B. (2017) Improved methods for multi-trait fine mapping of pleiotropic risk loci. Bioinforma. Oxf. Engl., 33, 248–255.

55. GTEx Consortium (2015) The Genotype-Tissue Expression (GTEx) pilot analysis: Multitissue gene regulation in humans. Science, 348, 648–660.

56. ENCODE Consortium (2004) The ENCODE (ENCyclopedia Of DNA Elements) Project. Science, 306, 636–640.

57. ENCODE Project Consortium (2012) An integrated encyclopedia of DNA elements in the human genome. Nature, 489, 57–74.

58. Wang, Y., Song, F., Zhang, B., Zhang, L., Xu, J., Kuang, D., Li, D., Choudhary, M.N.K., Li, Y., Hu, M., et al. (2018) The 3D Genome Browser: a web-based browser for visualizing 3D genome organization and long-range chromatin interactions. Genome Biol., 19, 151.

59. Nachman, M.W. and Crowell, S.L. (2000) Estimate of the mutation rate per nucleotide in humans. Genetics, 156, 297–304.

60. Zhao, Z. and Boerwinkle, E. (2002) Neighboring-Nucleotide Effects on Single Nucleotide Polymorphisms: A Study of 2.6 Million Polymorphisms Across the Human Genome. Genome Res., 12, 1679–1686.

61. Kitts, A., Phan, L., Ward, M. and Holmes, J.B. (2014) The Database of Short Genetic Variation (dbSNP) National Center for Biotechnology Information (US).

62. Niewold, T.B. (2015) Advances in Lupus Genetics. Curr. Opin. Rheumatol., 27, 440–447.

63. Patel, Z.H., Lu, X., Miller, D., Forney, C.R., Lee, J., Lynch, A., Schroeder, C., Parks, L., Magnusen, A.F., Chen, X., et al. (2018) A plausibly causal functional lupus-associated risk variant in the STAT1-STAT4 locus. Hum. Mol. Genet., 27, 2392–2404.

64. Parvin, J.D. and Sharp, P.A. (1993) DNA topology and a minimal set of basal factors for transcription by RNA polymerase II. Cell, 73, 533–540.

65. Scaffidi, P. and Bianchi, M.E. (2001) Spatially Precise DNA Bending Is an Essential Activity of the Sox2 Transcription Factor. J. Biol. Chem., 276, 47296–47302.

66. Kumasaka, N., Knights, A.J. and Gaffney, D.J. (2019) High-resolution genetic mapping of putative causal interactions between regions of open chromatin. Nat. Genet., 51, 128–137.

67. Yang, J., Fritsche, L.G., Zhou, X. and Abecasis, G. (2017) A Scalable Bayesian Method for Integrating Functional Information in Genome-wide Association Studies. Am. J. Hum. Genet., 101, 404–416.

68. Jiang, J., Cole, J.B., Freebern, E., Da, Y., VanRaden, P.M. and Ma, L. (2019) Functional annotation and Bayesian fine-mapping reveals candidate genes for important agronomic traits in Holstein bulls. Commun. Biol., 2, 212.

