## Supplemental Figures and Tables for "Intrinsic DNA topology as a prioritization metric in genomic fine-mapping studies"

**Figure S1. SKAT and SKAT-O analyses in *FAM167A-BLK*.**

Comparison of results for the *FAM167A-BLK* region using SKAT in a ΔMGW-weighted and equally weighted analysis. SNPs prioritized in the unweighted analyses (blue diamonds) were those that had multiple SNPs from the highly-associated LD block (Figure 5) within the same 5-SNP analysis windows (Table S3). Weighted analysis (shown in purple) instead prioritized SNPs that had larger ΔMGW and association values. This shifted the signal upstream to rs2061831. We note that the SKAT-O analysis performed very similarly to SKAT.

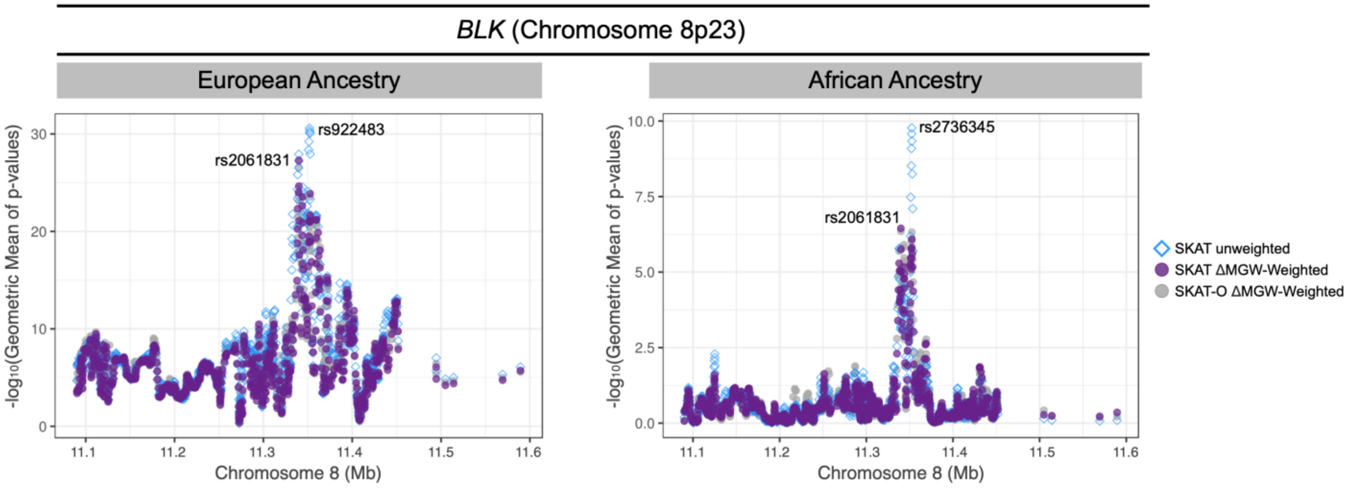

**Figure S2. Chromatin interactions (Hi-C) across cell types for identified SNPs in *FAM167A-BLK*.**

Comparison of Hi-C interactions for three SNPs in the *BLK-FAM167A* region. Data was queried using the 3D genome browser (http://promoter.bx.psu.edu/hi-c/). Each plot is centered on the labeled SNP with +/- 500 kb. Each arc indicates a chromatin interaction, as observed from Hi-C data. Chromatin interactions are only shown for those that have either a start or stop region that overlaps with the labeled SNP. The number of chromatin interactions with a SNP's region are indicated in parentheses for each plot. A majority of the observed chromatin interactions occur within 500 kb, with only a few extending further than the plotted region (e.g. rs2061831 for B-Cells, CD4-Cells, CD8-Cells, and Monocytes. For rs2736340, no chromatin interactions were observed in CD4-cells, CD8-cells, and Neutrophils. rs2061831 was strongly prioritized across ΔMGW-weighted analyses in both European and African Ancestries; and this SNP shows the most interactions for this data. The other two SNPs were identified in single-ancestry association analyses. Rs13277113 was the top SNP in EA while rs2736340 was the top SNP in AA analyses. Although all three SNPs are in high LD, they are physically separated from one another, eliciting different patterns of chromatin interactions.

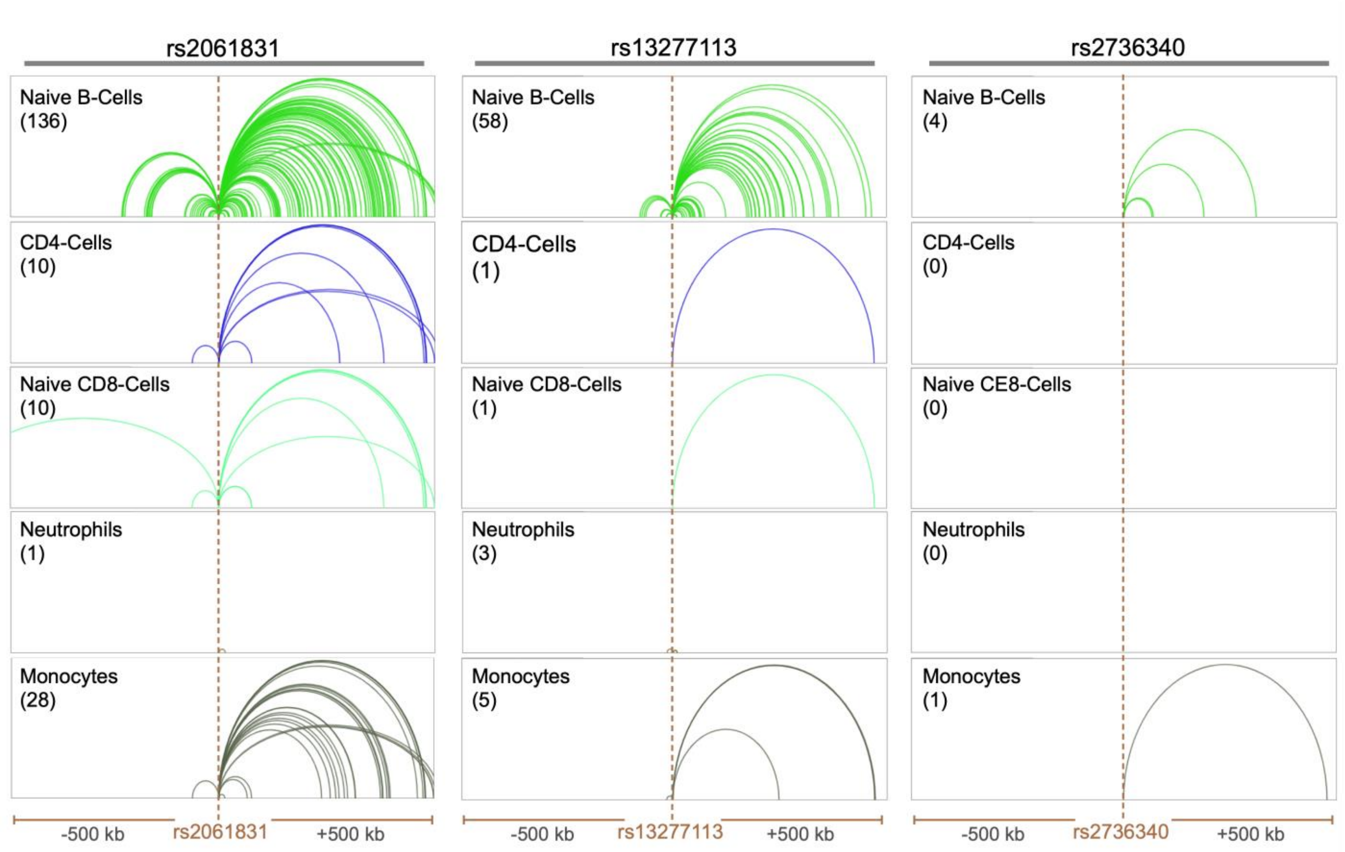

**Figure S3. SKAT and SKAT-O analyses in *STAT4*.**

Comparison of results for the *STAT4* region using SKAT in a ΔMGW-weighted (purple dots) and equally weighted analysis (blue diamonds). We note that the SKAT-O analysis performed very similarly to SKAT

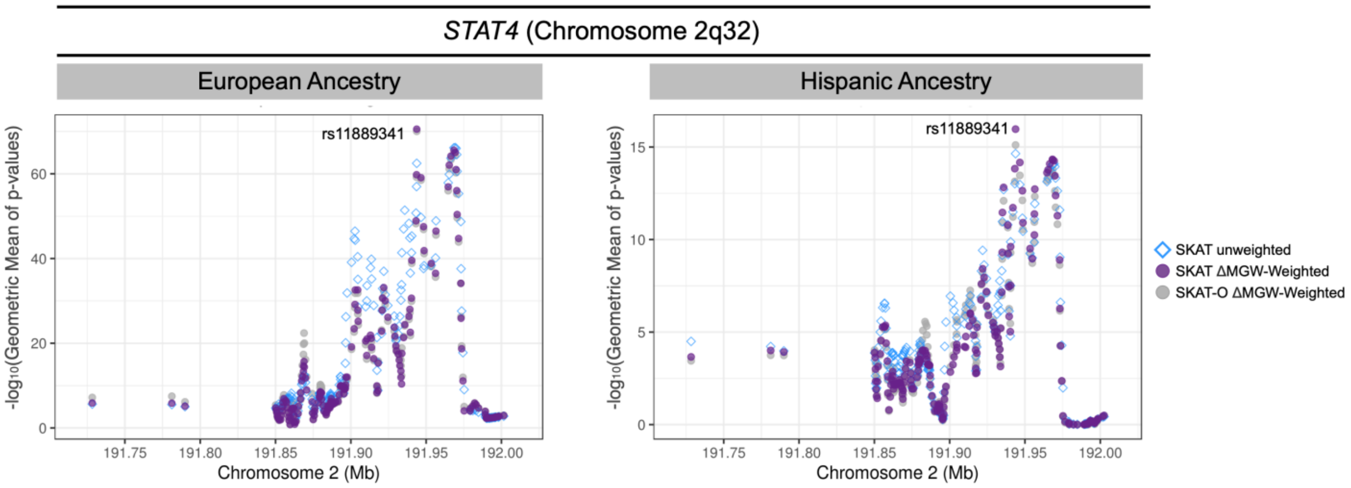

**Figure S4. SKAT and SKAT-O analyses in *TNIP1*.**

Comparison of results for the *TNIP1* region using SKAT in a ΔMGW-weighted and equally weighted analysis. Here, ΔMGW did not distinguish SNPs differently from the unweighted-analysis, other than an overall diminished prioritization signal, which is consistent for the low magnitudes of ΔMGW observed for the top-associated SNPs in the single-logistic regression analyses (Table S10-S11). In this region, SLE association, not ΔMGW, was the driver for prioritizing SNPs. We note that the SKAT-O analysis performed very similarly to SKAT

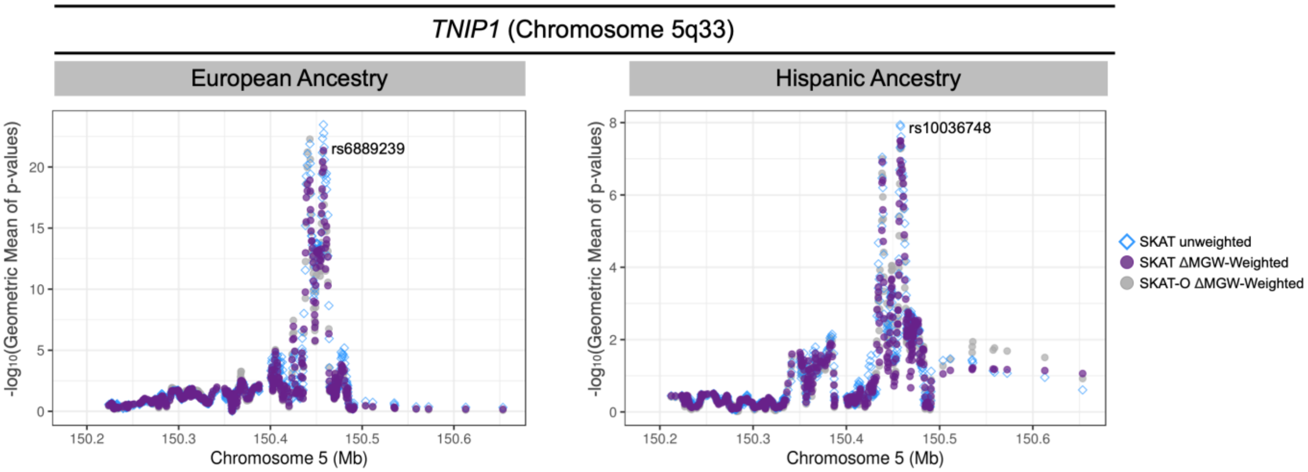

**Table S1. SNP counts for bi-allelic SNPs from dbSNP dataset 150.**

| SNP | SNP Count[a] | Percentage | Mutation Type | Count[b] | Percentage |
|---|---|---|---|---|---|
| A/G | 66,137,478 | 33.23 | Transition | 132,214,444 | 66.43 |
| C/T | 66,076,966 | 33.20 | | | |
| A/C | 17,265,950 | 8.67 | Transversion | 66,823,753 | 33.57 |
| A/T | 14,469,978 | 7.27 | | | |
| C/G | 17,783,460 | 8.93 | | | |
| G/T | 17,304,365 | 8.69 | | | |
| All | 199,038,197 | 100 | | 199,038,197 | 100 |

[a]Count bi-allelic SNPs that passed quality control filtering as stated in Methods. This excludes 75 SNPs that were not used for analysis due to uncertainty in flanking sequence (N in sequence).

[b]Cumulative sum for SNP Count, by Transition or Transversion mutation type. As established in the literature, transition mutations are more likely to occur, and here we observe 66.43% of included SNPs as transition mutations.

**Table S2. Summary of available NCBI SNP function designations.** Definitions are based on the NCBI Handbook, second edition. Counts are derived from SNP150 dataset.

| | | | Bi-allelic SNPs[a] with Function Label | |
|---|---|---|---|---|
| NCBI Function | Description | Sequence specificity | All -not exclusive[b] | Exclusive[c] |
| coding-synonymous | synonymous variant, the amino acid is not changed by either allele. | Yes. The change in the codon must yield same amino acid. | 1,863,763 | 1,178,980 |
| intron | variant within non-coding region of a gene. | No | 89,334,045 | 84,909,115 |
| missense | variant alters codon to change amino acid product. | Yes. The change in the codon must yield different amino acid. | 3,723,326 | 2,345,831 |
| ncRNA | variant within a non-coding RNA | No | 2,266,252 | 499,593 |
| near-gene-3 | located within 500 bases of a gene, 3'. | No | 1,315,373 | 654,589 |
| near-gene-5 | located within 2000 bases of a gene, 5'. | No | 5,418,679 | 2,487,192 |
| nonsense | variant produces a stop codon. | Yes. Variant must produce stop Codon (TGA) | 112,826 | 66,275 |
| splice-3 | variant located in the last two bases in the 3' end of an intron. | Most introns end with AG | 37,092 | 25,401 |
| splice-5 | variant located in the first two bases in the 5' end of an intron. | Most introns start with GU | 43,940 | 28,983 |
| stop-loss | variant changes STOP codon to non-stop codon. | Yes. Variant allele must produce a codon that is not a stop codon | 5,357 | 2,225 |
| unknown | variant not within other categories. Intergenic regions. | No | 99,004,130 | 99,004,130 |
| untranslated-3 | located within 3' untranslated region | No | 2,255,433 | 1,299,685 |
| untranslated-5 | located within 5' untranslated region. | No | 922,224 | 181,208 |

[a] Data from the dbSNP SNP150 dataset. Counts refer to post-quality control pruning as described in Methods. This includes exclusion of 75 SNPs which had uncertainty in flanking sequence.
[b] SNPs can have more than one label. A SNP may be labeled as both as ‘near-gene-3’ and ‘ncRNA’. Thus, a single SNP can contribute to more than one functional category count.
[c] SNPs with only one NCBI designated function (i.e. intron, only). This represents a subset of the SNPs counted in the “all-not exclusive” variable. The exception is for the “unknown SNPs” category. All designated “unknown” SNPs are designated because they do not fall within any other categories. Hence, here, the exclusive and not-exclusive counts are the same.

**Table S3**. **Summary of ΔMGW measures in the *FAM1167A-BLK* Region**

| | 500 kb Region | | 60 kb region[a] | |
|---|---|---|---|---|
| | EA | AA | EA | AA |
| N SNPs[b] | 835 | 933 | 145 | 159 |
| Mean ΔMGW (Å) | 0.62 | 0.63 | 0.62 | 0.61 |
| Median ΔMGW (Å) | 0.53 | 0.53 | 0.49 | 0.50 |
| Minimum ΔMGW (Å) | 0.07 | 0.07 | 0.12 | 0.12 |
| Maximum ΔMGW (Å) | 2.72 | 3.16 | 1.97 | 3.16 |
| σ[c] | 84 | 103 | 15 | 17 |
| 2σ[d] | 21 | 28 | 3 | 4 |

[a]Genomic region encompassing primary peak of association, as shown in Figure 5
[b]Number of SNPs that passed quality control measures. Within a single population, SNPs that are monomorphic are excluded.
[c]Number of SNPs greater than one Standard Deviation (>1.11 Å) from the Mean ΔMGW in the NCBI SNP150 data
[d]Number of SNPs greater than two Standard Deviations (> 1.54 Å) from the Mean ΔMGW in the NCBI SNP150 data

**Table S4. EA results in the 60kb association peak in FAM167A-BLK region.**

*Excel File.*

**Table S5. AA results in the 60kb association peak in FAM167A-BLK region.**

*Excel File.*

**Table S6**. **Summary of ΔMGW measures in the *STAT4* Region**

| | 500 kb Region | | 11 Mb region[a] | |
|---|---|---|---|---|
| | EA | HA | EA | HA |
| N SNPs[b] | 192 | 202 | 111 | 118 |
| Mean ΔMGW (Å) | 0.73 | 0.74 | 0.79 | 0.78 |
| Median ΔMGW (Å) | 0.57 | 0.57 | 0.58 | 0.58 |
| Minimum ΔMGW (Å) | 0.11 | 0.11 | 0.11 | 0.11 |
| Maximum ΔMGW (Å) | 3.16 | 3.16 | 3.16 | 3.16 |
| σ[c] | 31 | 36 | 22 | 23 |
| 2σ[d] | 15 | 16 | 12 | 12 |

[a]Genomic region encompassing primary peak of association, as shown in Figure 6

[b]Number of SNPs that passed quality control measures. Within a single population, SNPs that are monomorphic are excluded.

[c]Number of SNPs greater than one Standard Deviation (>1.11 Å) from the Mean ΔMGW in the NCBI SNP150 data

[d]Number of SNPs greater than two Standard Deviations (> 1.54 Å) from the Mean ΔMGW in the NCBI SNP150 data

**Table S7. EA results in the 11Mb association peak in *STAT4* region.**

*Excel File.*

**Table S8. HA results in the 11Mb association peak in *STAT4* region.**

*Excel File.*

**Table S9. Summary of ΔMGW measures in the *TNIP1* Region**

| | 500 kb Region | | 40 kb region[a] | |
|---|---|---|---|---|
| | EA | HA | EA | HA |
| N SNPs[b] | 496 | 500 | 99 | 103 |
| Mean ΔMGW (Å) | 0.67 | 0.67 | 0.61 | 0.62 |
| Median ΔMGW (Å) | 0.55 | 0.55 | 0.51 | 0.52 |
| Minimum ΔMGW (Å) | 0.11 | 0.11 | 0.11 | 0.11 |
| Maximum ΔMGW (Å) | 2.31 | 2.31 | 2.29 | 2.29 |
| σ[c] | 67 | 66 | 10 | 11 |
| 2σ[d] | 23 | 24 | 4 | 4 |

[a]Genomic region encompassing primary peak of association, as shown in Figure 7

[b]Number of SNPs that passed quality control measures. Within a single population, SNPs that are monomorphic are excluded.

[c]Number of SNPs greater than one Standard Deviation (>1.11 Å) from the Mean ΔMGW in the NCBI SNP150 data

[d]Number of SNPs greater than two Standard Deviations (> 1.54 Å) from the Mean ΔMGW in the NCBI SNP150 data

**Table S10. EA results in the 40kb association peak in *TNIP1* region.**

*Excel File.*

**Table S11. HA results in the 40kb association peak in *TNIP1* region.**

*Excel File.*
